# Iterative *in vivo* cut’n’paste of chromosomal loci in *Escherichia coli* K-12 using synthetic DNA

**DOI:** 10.1101/2025.06.19.660532

**Authors:** Sebastian Gude, Andreas B. Bertelsen, Morten H. H. Nørholm

**Author notes:** ^2^Novo Nordisk A/S, Hørsholm, Denmark (Morten H. H. Nørholm). **Corresponding author:** Sebastian Gude, The Novo Nordisk Foundation Center for Biosustainability, Technical University of Denmark, Kongens Lyngby, Denmark.

## Abstract

We have developed an easy-to-use *in vivo* method to iteratively relocate functional chromosomal loci onto an episome in *Escherichia coli* by utilizing synthetic DNA fragments. In this *in vivo* cut’n’paste procedure, “cutting” is executed by the RNA-guided DNA endonuclease Cas9 and a set of guides, while “pasting” is facilitated by the phage λ Red recombinase, which are all synthesized on easily curable helper plasmids. To demonstrate the utility of *in vivo* cut’n’paste, we commercially obtained synthetic DNA fragments containing locus-specific homology regions, antibiotic marker cassettes, and standardized Cas9 target sequences, and successfully relocated seven functional chromosomal loci. Scarless relocation mutants of the *trg*, *aer*, *tsr*, *malHM*, *malQT*, *macB*-*nadA*, and *folA* chromosomal loci were obtained as antibiotic-resistant isolates by combining Cas9 counterselection with the restoration of an antibiotic marker cassette. The additional antibiotic marker cassettes and standardized Cas9 target sequences present in the synthetic DNA fragments are inherently eliminated upon completion of the procedure, enabling iterative processing of chromosomal loci. *In vivo* cut’n’paste should be widely useful, especially in genome (re-)engineering efforts of *E. coli* and other bacteria because the procedure can be performed by non-specialists in unmodified wild-type cells.

**Significance:** Editing of genomes is crucial to understand the inner workings of bacteria and to alter them for biotechnological applications. *In vivo* cut’n’paste, in analogy to the frequently used keyboard shortcuts Ctrl-X and Ctrl-V which allow to move text, images, or files on computers, makes it easy to relocate large chunks of the bacterial genome without requiring skills to manipulate DNA outside of the bacterial cell. Like Ctrl-X and Ctrl-V, *in vivo* cut’n’paste can be executed several times to iteratively edit multiple genomic locations in the same bacterium. Since *in vivo* cut’n’paste entirely takes place inside the bacterial cell, it is particularly well suited to edit hard-to-tackle genomic features such as essential genes.

## Introduction

Democratising access to methodological advancements, i.e., making them accessible to a broad audience of non-specialists, is a key component of method development. The introduction of the point-and-click graphical user interface by Apple and Microsoft^1^, the creation of the “batteries included” programming language Python^2^, and the development of the one-step inactivation of chromosomal genes procedure^3^ are a few prominent examples of such endeavours. Graphical user interfaces remove the hurdle to know how to interact with a computer via a text console. The extensive Python Standard Library supports nearly every aspect of computer science, thus, freeing programmers from having to reimplement many commonly required features over and over again. Similarly, the one-step inactivation of chromosomal genes procedure enables researchers to generate gene knockouts without the need to assemble integrative plasmids.

The ability to manipulate genomes is key for uncovering and leveraging the potential of (micro-)biology. A milestone in our ability to edit bacterial genomes was the development of the one-step inactivation of chromosomal genes procedure^3^. Among many other scientific and engineering endeavours, it facilitated the creation of a comprehensive single gene knockout library and the charting of essential genes in *E. coli*^4^, the study of bacterial motility^5,6^, stringent response^7^, the interplay of growth and gene expression^8^, two-component systems^9^, evolution of antibiotic resistsance^10^, and chromosomal organisation^11^ as well as the development of strains with reduced genomes^12^ and highly titratable induction systems^13^ while also playing a crucial role in metabolic engineering applications^14,15^. Though the one-step inactivation procedure^3^ was preceded by numerous other bacterial genome editing techniques^16–23^, its ease of use and broad applicability ensured its still continuing success. The required short homology extensions to create genomic deletions are easily attached to an antibiotic marker cassette via PCR, the recombineering machinery is expressed from an easily curable plasmid, and the procedure does not require any prior modifications to the bacterial genome^3^. Furthermore, the antibiotic marker cassette can subsequently be eliminated via FLP-FRT recombination, resulting in an unmarked, yet not fully scarless, mutant that is compatible with another round of genome editing^3^.

With the rise of CRISPR-Cas technologies^24^, λ Red recombineering was successfully supplemented with sequence-specific RNA-guided counterselection^25–27^. Combining λ Red recombineering and CRISPR-Cas with intricate multi-layered selection, counterselection, and colour screening enables large-scale genome editing such as genome recoding^28,29^, global genome rearrangement^30,31^, and interspecies gene transfer^32^. While these sophisticated large-scale CRISPR-Cas genome editing techniques accomplish feats vastly beyond the capabilities of the one-step inactivation of chromosomal genes procedure^3^, they unfortunately fall short in one aspect. Namely, they are relatively difficult to implement in non-specialist laboratories due to their technical complexity, thus impeding the democratisation of large-scale genome editing.

Building upon established large-scale CRISPR-Cas genome editing methods^28–32^, we present here *in vivo* cut’n’paste - an attempt to democratise large-scale genome editing. *In vivo* cut’n’paste is an easy-to-use procedure to relocate chromosomal loci onto an episome in wild-type cells that was designed to be performed by non-specialists without expert skills in molecular biology. *In vivo* cut’n’paste solely relies on commercially synthesized DNA fragments and a set of easily curable helper plasmids. Leveraging CRISPR-Cas cutting and λ Red recombineering with commercially synthesized DNA fragments completely removes the need for *in vitro* DNA manipulations. Furthermore, easily curable helper plasmids and an antibiotic marker recycling scheme inherently enable the iterative creation of multiple large-scale genome edits. We successfully relocated seven functional genomic loci, including metabolic and essential genes, in *E. coli* of up to 140 kb in size.

## Results

### *In vivo* cut’n’paste

*In vivo* cut’n’paste enables to iteratively relocate genomic loci from the native chromosome onto an episome (Fig. 1A, Supp. Movie 1). In each round, a genomic locus, defined by two pairs of tandem homology regions HR^A^-HR^B^ and HR^C^-HR^D^ is excised from the native chromosome and subsequently integrated in between the homology regions HR^E^ and HR^TET^ on an episome (Fig. 1B). The entire *in vivo* cut’n’paste procedure occurs inside the cell without requiring any *in vitro* DNA manipulations.

**Figure 1:**
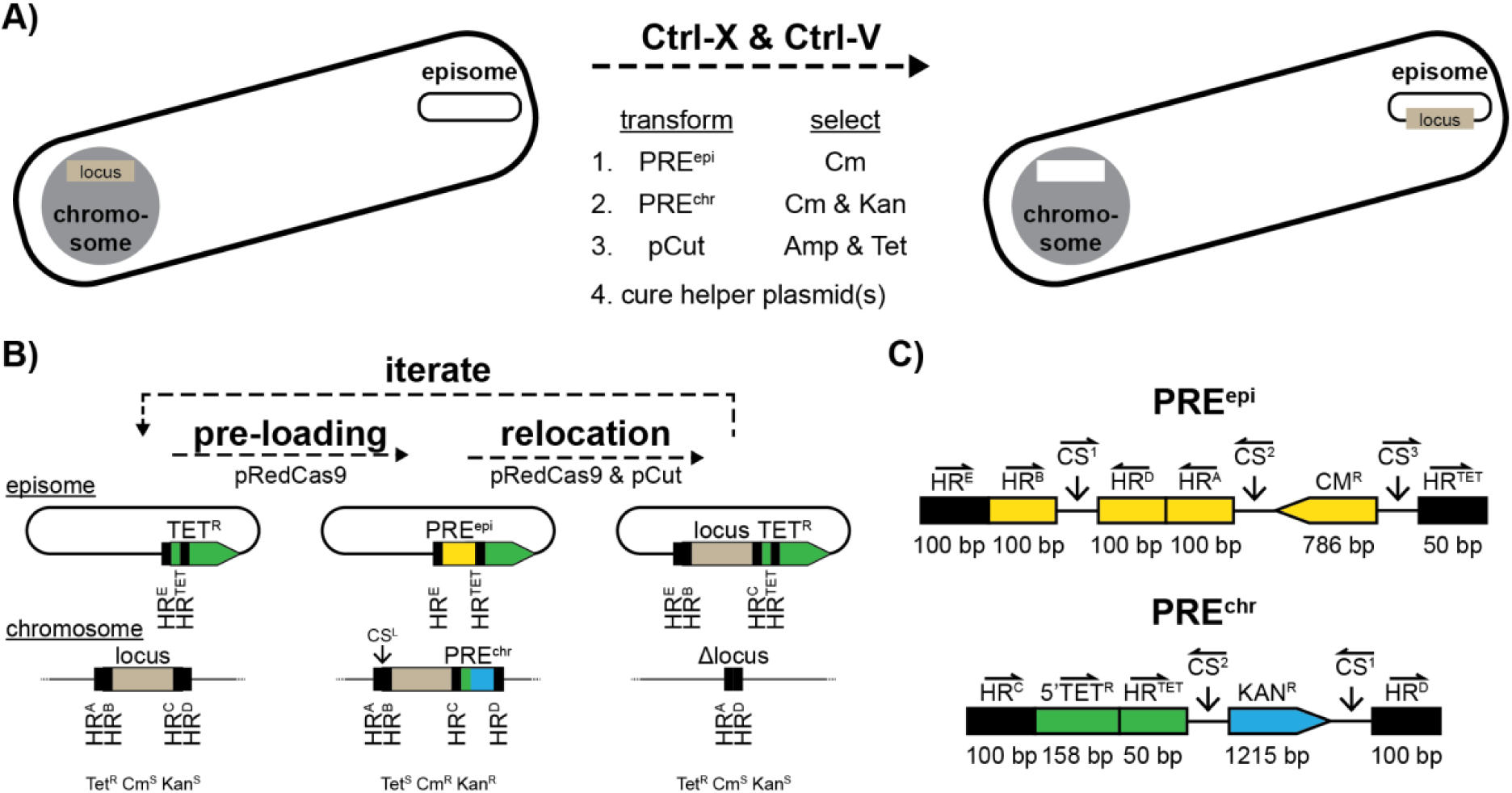
Iterative *in vivo* cut’n’paste. **a)** Graphical overview of *in vivo* cut’n’paste. **b)** Outline of the individual steps (pre-loading and relocation) of the *in vivo* cut’n’paste procedure. **c)** Features of commercially synthesized DNA fragments PRE^epi^ and PRE^chr^. A more detailed description of the design choices is provided in the Supplementary Materials. Half-arrows indicate the relative orientations of features. Solid, downwards arrows mark constant cut sites. pRedCas9 and pCut: easily curable helper plasmids, HR: homology region, CS: cut site, TET^R^: tetracycline marker cassette, CM^R^: chloramphenicol marker cassette, KAN^R^: kanamycin marker cassette, 5’TET^R^: non-functional fragment of TET^R^, Tet: tetracycline, Cm: chloramphenicol, Kan: kanamycin, R: resistant, S: susceptible, and bp: base pairs. Not to scale.

Each round of *in vivo* cut’n’paste consists of two phases: (1) pre-loading and (2) relocation (Fig. 1B). During pre-loading, two commercially synthesized DNA fragments, PRE^epi^ and PRE^chr^, are sequentially integrated via homologous recombination into the episome and the native chromosome, respectively, to equip cells with the required modification for the subsequent relocation of the genomic locus (Fig. 1C; see Supplementary Materials and Supp. Fig. 1-3 for a detailed description of pre-loading fragment design choices). Both pre-loading events are accompanied by positive antibiotic selection.

PRE^epi^ is flanked by the homology regions HR^E^ and HR^TET^. Directly downstream of HR^E^, PRE^epi^ contains the inside homology region HR^B^ of the genomic locus. The two outside homology regions HR^A^ and HR^D^ of the genomic locus are located next to HR^B^ followed by a chloramphenicol marker cassette (Fig. 1C). Integration of PRE^epi^ at HR^E^ and HR^TET^ into the episome truncates the tetracycline maker of the episome and, thus, renders the cells sensitive to tetracycline yet resistant to chloramphenicol (Fig. 1B).

PRE^chr^ is flanked by the homology region HR^C^ and HR^D^ of the genomic locus, enabling its integration directly adjacent to the genomic locus of interest in the native chromosome (Fig. 1B). In PRE^chr^, HR^C^ is directly followed by 5’TET^R^, encoding a short, non-functional fragment of the tetracycline marker cassette, and HR^TET^. A kanamycin marker cassette completes PRE^chr^ (Fig. 1C). Notably, the pre-loading fragments can be commercially obtained in their entirety without requiring further assembly.

Once pre-loading is completed, cells contain all features required for relocation (Fig. 1B,C). Firstly, the genomic locus in the native chromosome now contains HR^B^ on one side and HR^TET^ on its other side. Secondly, the episome also contains both HR^B^ and HR^TET^. Thirdly, a “repair band-aid”, consisting of HR^A^-HR^D^, which can fix the double-stranded break that will be created by the excision of the genomic locus from the native chromosome, is present on the episome (see Supp. Movie 1 for a detailed, step-by-step illustration of the pre-loading process). Optionally, additional features may be introduced during pre-loading. For example, a terminator can be inserted in between HR^C^ and 5’TET^R^ in EPI^chr^, which will, once relocation is completed, partially insulate the relocated locus from its new local environment on the episome.

Relocation is initiated by transforming the helper plasmid pCut (Fig. 1B). pCut constitutively expresses standardized, functionally tested guides targeting constant cut sites (CS^1^, CS^2^, and CS^3^) included in the pre-loading fragments from a CRISPR array, as well as a single guide (CS^L^) targeting a locus-specific junction between HR^A^ and HR^B^ (Fig. 1B,C). The sequence encoding the locus-specific single guide can effectively be inserted as a third synthetic DNA fragment into the helper plasmid pCut via I-CreI cutting counterselection-aided plasmid recombineering, thus completely eliminating the need for *in vitro* DNA manipulations (see Supp. Fig. 4 for details regarding I-CreI cutting counterselection-aided plasmid recombineering; a technical description of the procedure is provided in the Methods). In total, RNA-guided Cas9 endonuclease activity now introduces double-stranded breaks into six targeted locations across the episome and the native chromosome (Fig. 1B,C). These double-stranded breaks release the genomic locus from the native chromosome and the “repair band-aid” from the episome while simultaneously exposing the required homology regions for integration of the genomic locus into the episome and for integration of the “repair band-aid” into the native chromosome. Importantly, integration of the genomic locus into the episome not only fixes its double-stranded break but, additionally, leads to the restoration of the tetracycline marker cassette, enabling positive selection of successful relocation events in the presence of tetracycline (see Supp. Movie 1 for a detailed step-by-step illustration of the relocation process).

The chloramphenicol and kanamycin marker cassettes, along with the constant cut sites, which were all introduced during pre-loading as components of PRE^epi^ and PRE^chr^, are inherently lost during the relocation process. Finally, the helper plasmid pCut encodes rhamnose-inducible, self-targeting guides enabling its straightforward elimination from cells (see Methods for details), thus immediately enabling relocation mutants to undergo additional rounds of *in vivo* cut’n’paste (Fig. 1B). We typically tested four isolates by diagnostic PCR at each editing step (episome pre-loading, native chromosome pre-loading, and relocation) of the *in vivo* cut’n’paste procedure. An individual isolate possessing the desired genomic alternation was chosen for further processing.

### Relocating a genomic locus

In a first attempt, we employed *in vivo* cut’n’paste to relocate the *trg* locus from its native genomic location in strain CNP^empty^ (*E. coli* K-12 MG1655 WT + EPI^empty^) onto an empty episome (EPI^empty^). Candidates of successful relocations, CNP^trg^ (*E. coli* K-12 MG1655 Δ*trg* + EPI^trg^), were obtained under tetracycline selection. After curing the helper plasmids, we sent a CNP^trg^ isolate for short-read whole genome re-sequencing (Supp. Table 1). Mapping reads of the CNP^trg^ isolate to the parental CNP^empty^ reference depicted a depth of coverage of about 400x across the native chromosome and the episome (Fig. 2A, middle row). No mutations relative to the parental strain CNP^empty^ were observed. Yet, notably, pairs of adjacent bases at three genomic locations were not spanned by any reads. Namely, both edges of the *trg* locus on the native chromosome (HR^A^-HR^B^ and HR^C^-HR^D^) and one edge of HR^E^ on the episome lacked spanning reads (Fig. 2A, middle row). No lack of spanning reads at these locations was observed when reads of the parental CNP^empty^ strain were mapped to the CNP^empty^ reference (Fig. 2A, top row). The lack of spanning reads in the CNP^trg^ isolate suggested that though the *trg* locus was present, it was not located in its native genomic location, and, additionally, that something had likely been inserted into the episome next to HR^E^. We updated the reference map to include the desired relocation of the *trg* locus based on our *in silico* design of the *in vivo* cut’n’paste procedure. When the reads of the CNP^trg^ isolate were mapped to this updated reference, the lack of spanning reads disappeared (Fig. 2A, bottom row), suggesting that the *trg* locus had indeed been relocated from the native chromosome onto the episome. Long-read sequencing of an extracted episome further supported the successful relocation of the *trg* locus (Supp. Table 2). In total, we observed eleven reads that spanned from the backbone of the episome across the entire, mutation-free *trg* locus.

**Figure 2:**
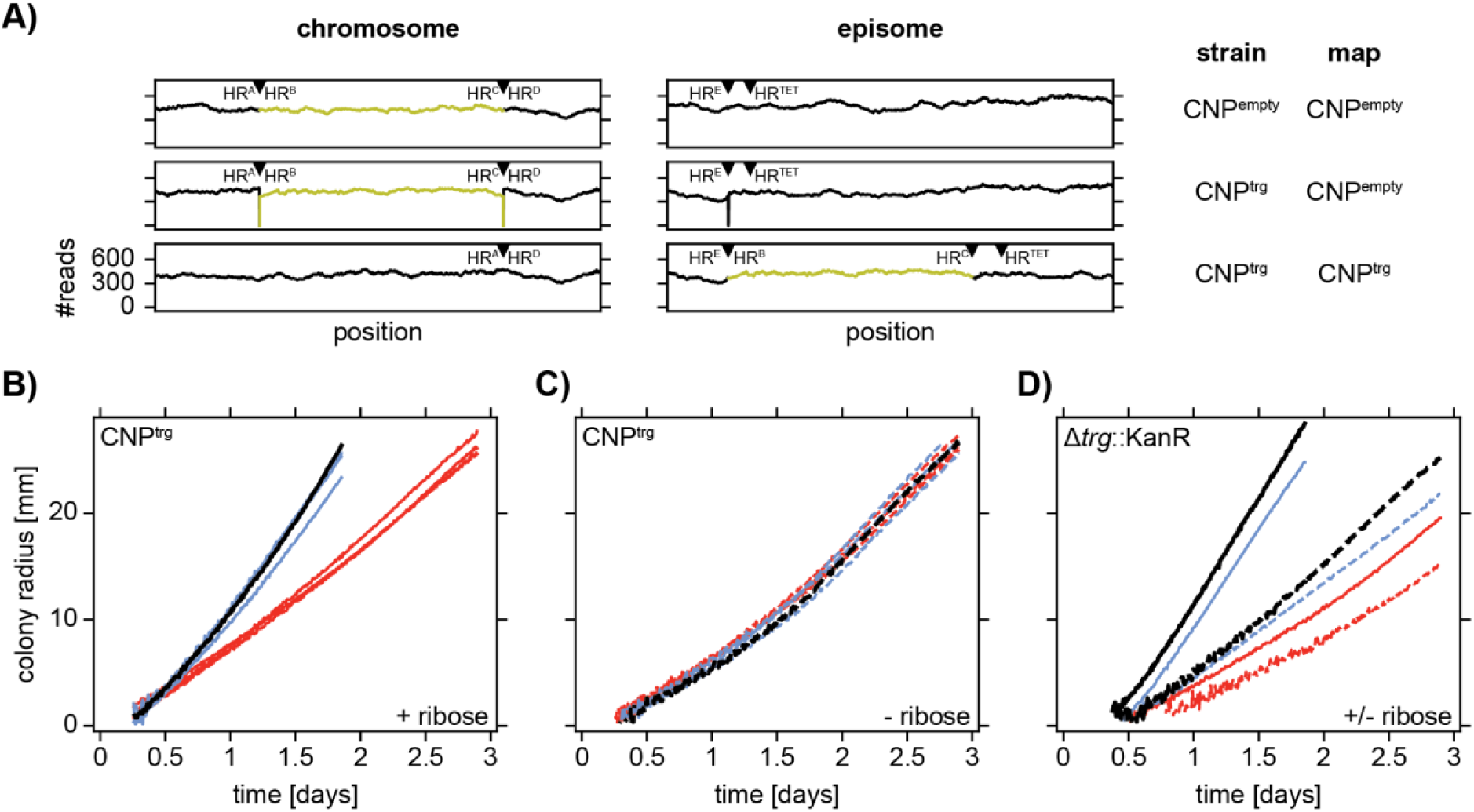
I*n vivo* cut’n’paste of the *trg* locus. **a)** Number of mapped reads in 3.2 kb segments of the native chromosome and the episome. Only reads mapping to a position and to both of its direct neighbors are counted. Strain identities and reference maps as indicated within the panel. The *trg* locus is highlighted in yellow. Triangles mark locations of homology regions. **b-d)** Motility phenotypes in soft agar. Colony expansion as function of time in glycerol minimal medium with (solid) or without (dashed) 100 μM D-ribose. **b-c)** CNP^trg^ (blue), CNP^trg^ without EPI^trg^ (red), and CNP^empty^ (black). Three independent CNP^trg^ isolates and a single replicate of CNP^empty^. **d)** *E. coli* K-12 MG1655 Δtrg::KanR with EPI^trg^ (blue), *E. coli* K-12 MG1655 Δtrg::KanR with EPI^empty^ (red), and CNP^empty^ (black). A single replicate each.

The *trg* locus (≍1.8 kb) contains a single gene encoding the transmembrane Trg chemoreceptor. Trg mediates directed motile behaviour (i.e., chemotaxis) toward D-ribose^33^. We assayed chemotactic colony expansion dynamics in glycerol minimal medium soft agar plates in the presence and absence of low supplementations of D-ribose as proxy for ribose sensing (see Methods for details). Initially, a small drop of bacterial culture was spotted into the centre of the soft agar plates. Upon growing, bacteria locally deplete nutrients and thus create gradients enabling chemotactic sensing and expansion^34^. Expansion dynamics of *trg*-relocated CNP^trg^ isolates (blue) in glycerol minimal medium soft agar plates in the presence (sold lines) and absence (dashed lines) of D-ribose were comparable to the behaviour of their ancestor CNP^empty^ (black), clearly indicating that *trg* retained its functionality after relocation onto the episome (Fig. 2B,C). When the episome EPI^trg^ was removed from the CNP^trg^ isolates (red), the expansion dynamics in presence of D-ribose were reduced to a slower aerotactic behaviour^35^ as also observed in glycerol minimal medium in the absence of D-ribose, indicating that the ability to sense D-ribose and utilizing its concentration gradients for chemotactic expansion was lost along with the episome (Fig. 2B,C). As an additional test, we separately transferred the episomes EPI^trg^ and EPI^empty^ into a strain background in which the *trg* gene had been disrupted by a kanamycin marker cassette (*E. coli* K-12 MG1655 Δtrg::KanR). As expected, faster chemotactic expansion dynamics in presence of D-ribose were restored only when EPI^trg^, containing the relocated *trg* locus, was present (Fig. 2D).

### Iteratively relocating genomic loci

In accordance with the design principles of the *in vivo* cut’n’paste procedure, the additional antibiotic marker cassettes and constant cut sites acquired during pre-loading were inherently lost, and the tetracycline marker cassette on the episome was restored, upon completion of the *trg* relocation (Fig. 1B). Thus, CNP^trg^ was capable of undergoing further rounds of *in vivo* cut’n’paste. We iteratively applied *in vivo* cut’n’paste to relocate two additional genomic loci onto the episome using CNP^trg^ as a starting point (Fig. 3A). First, the *aer* locus (≍1.7 kb) encoding the aerotaxis receptor Aer was relocated, resulting in CNP^trg,aer^ (*E. coli* K-12 MG1655 Δ*trg* Δ*aer* + EPI^trg,aer^). Then, the *tsr* locus (≍1.8 kb) encoding the chemoreceptor Tsr, mediating aerotaxis and sensing of L-Serine, was relocated, resulting in CNP^trg,aer,tsr^ (*E. coli* K-12 MG1655 Δ*trg* Δ*aer* Δ*tsr* + EPI^trg,aer,tsr^).

**Figure 3:**
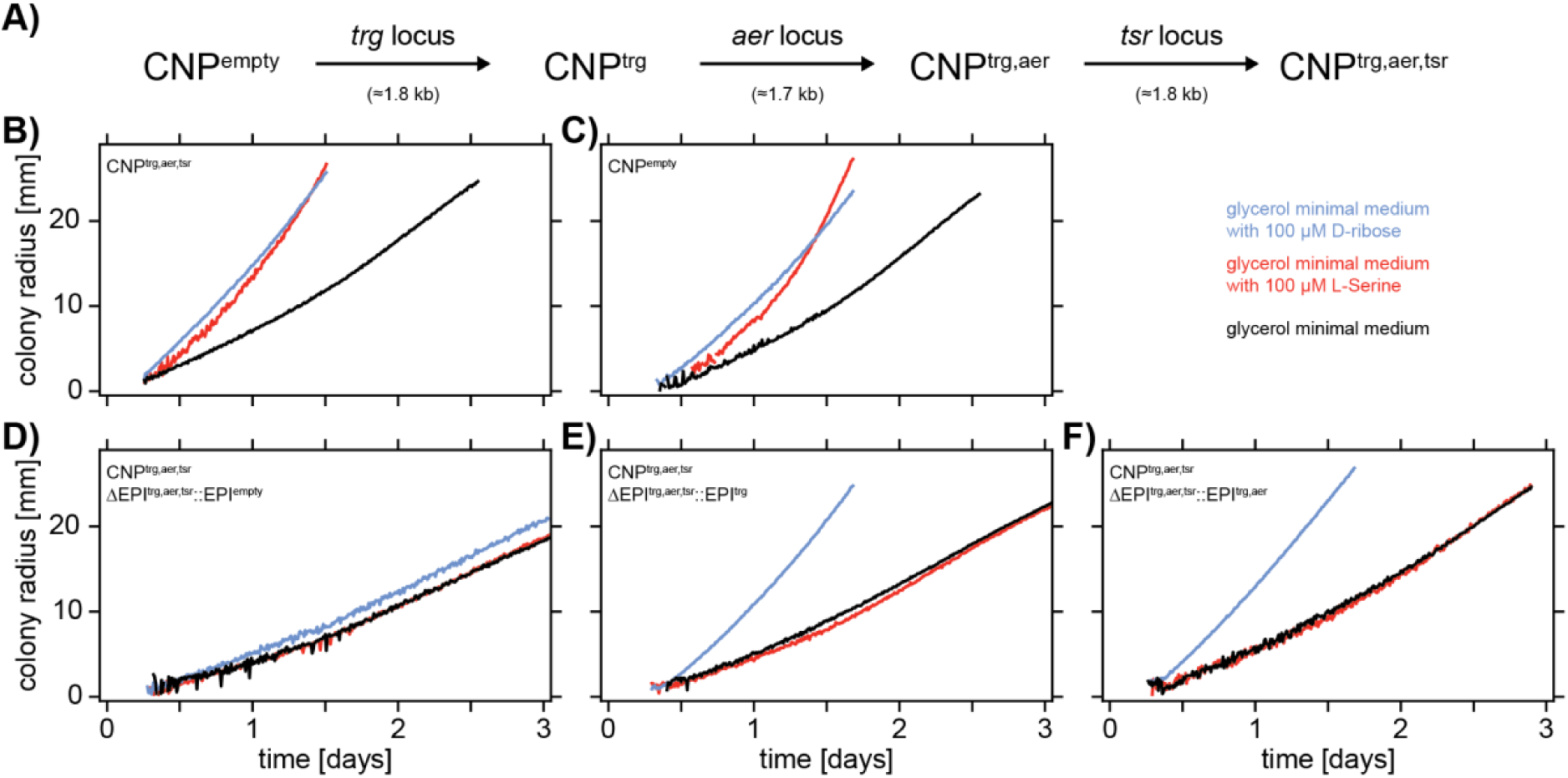
Iterative *in vivo* cut’n’paste of three chemoreceptor loci. **a)** Illustration of iterative *in vivo* cut’n’paste of the *trg*, *aer*, and *tsr* chemoreceptor loci. **b-f)** Motility phenotypes in soft agar. Colony expansion as function of time in glycerol minimal medium with 100 μM D-ribose (blue), with 100 μM L-Serine (red), or without any additions (black). Strain identities as defined within the individual panels.

The relocated loci were mutation-free (Supp. Table 2). The triple-relocated CNP^trg,aer,tsr^ qualitatively retained the colony expansion behaviour of its ancestor CNP^empty^ in presence of D-ribose, L-Serine, or in the absence of supplementations to glycerol minimal medium (Fig. 3 B,C). Furthermore, separately transferring the episomes EPI^empty^ (Fig. 3D), EPI^trg^ (Fig. 3 E), and EPI^trg,aer^ (Fig. 3F) into a Δ*trg* Δ*aer* Δ*tsr* chromosomal background (i.e., a CNP^trg,aer,tsr^ derivative from which EPI^trg,aer,tsr^ had been eliminated) partially restored the colony expansion behaviour in presence of D-ribose (sensed via Trg), L-Serine (sensed via Tsr), and in absence of any supplementations (energytaxis via Aer and Tsr) as expected^33,35^.

In an independent attempt, using CNP^empty^ as a starting point, we successfully iteratively relocated two genomic loci (≍8.1 kb and ≍7.9 kb) encoding 10 genes involved in maltose utilisation onto an episome (Fig. 4A). Again, the relocated loci were mutation-free. The resulting relocation mutants CNP^malHM^ (*E. coli* K-12 MG1655 Δ(*malH*-*malM*) + EPI^malHM^) and CNP^malHM,malQT^ (*E. coli* K-12 MG1655 Δ(*malH*-*malM*) Δ(*malQ*-*malT*) + EPI^malHM,malQT^) retained their ability to uptake and ferment maltose as indicated by producing a dark red phenotype when spotted and grown on maltose-MacConkey/agar plates (Fig. 4B).

**Figure 4:**
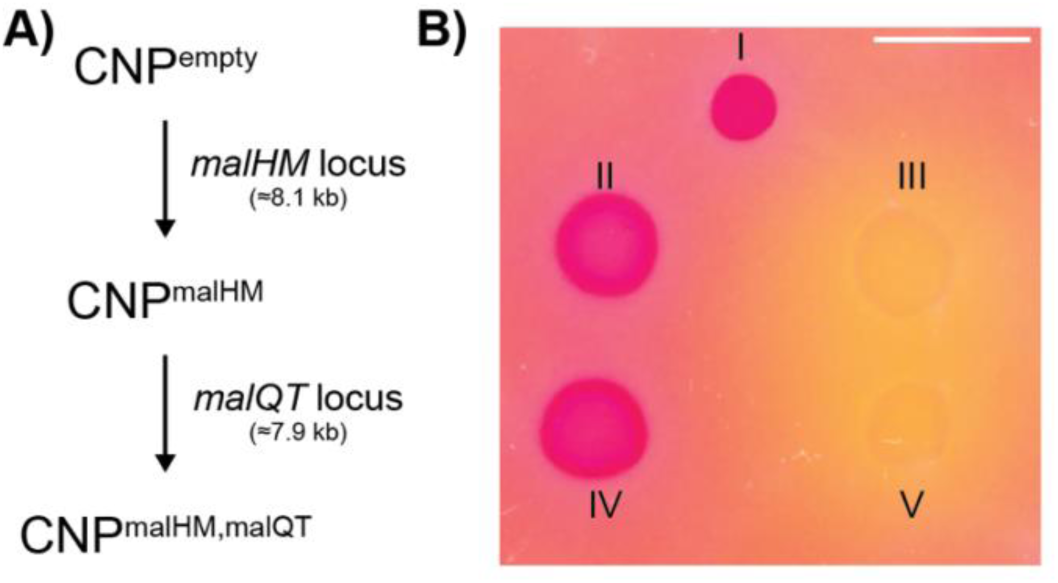
Iterative in vivo cut’n’paste of two maltose-utilization loci. **a)** Illustration of iterative *in vivo* cut’n’paste of the *malHM* (i.e., *malEFGH* and *malKlamBmalM*) and *malQT* (i.e., *malPQ* and *malT*) maltose-utilization loci. **b)** Maltose utilization of CNP strains. Spotting of liquid culture onto maltose-MacConkey+Tet/agar. Imaged after 9 hours. Dark red disks: cells can utilize maltose, pale yellow disks with halo: cells cannot utilize maltose. The orange-reddish background depicts MacConkey/agar. I: CNP^empty^, II: CNP^malHM^, III: CNP^malHM^ ΔEPI^malHM^::EPI^empty^, IV: CNP^malHM,malQT^, and V: CNP^malHM,malQT^ ΔEPI^malHM,malQT^::EPI^empty^. Scale bar: 10 mm.

### Relocating a large genomic locus and an essential gene

To illustrate the utility of *in vivo* cut’n’paste to process large genomic loci, we attempted to relocate the *macB*-*nadA* locus (≍140 kb) containing 135 non-essential genes from its native genomic location onto an episome. The relocation was successful and mutation-free (Supp. Table 1).

As opposed to traditional sequential deletion and complementation procedures, all genes, including those located inside the genomic locus that is edited, remain present throughout the entire *in vivo* cut’n’paste process. Hence, *in vivo* cut’n’paste should be well suited for manipulating hard-to-edit genomic features such as essential genes. Indeed, we successfully relocated the essential gene *folA* onto an episome, resulting in CNP^folA^ (*E. coli* K-12 MG1655 Δ*folA* + EPI^folA^).

An episome containing an essential gene that is simultaneously absent from the native chromosome is expected to be stably maintained even in the absence of antibiotic selection because cells are unable to survive without the essential gene. As a direct test, we removed the tetracycline antibiotic marker cassette from EPI^folA^ and inserted a highly expressed green fluorescent reporter (GFP) in its place, resulting in CNP^folA,TetR::gfp^ (*E. coli* K-12 MG1655 Δ*folA* + EPI^folA,TetR::gfp^). We then assayed the retention of the antibiotic marker-free episome EPI^folA,TetR::gfp^ by serial culturing of CNP^folA,TetR::gfp^ in LB medium without any antibiotic supplementation (see Methods for details). Plating on non-selective LB agar plates was performed at each transfer. After overnight incubation, plates were imaged in GFP and brightfield channels (Fig. 5, Supp. Fig. 5). At each transfer, all fluorescent colonies visible in the GFP channel (Fig. 5A) had a corresponding colony in the associated brightfield image (Fig. 5B) and vice versa. Thus, 100% of the bacterial population stably retained the antibiotic marker-free episome EPI^folA,TetR::gfp^ throughout ten serial passages in the absence of antibiotic selection.

**Figure 5:**
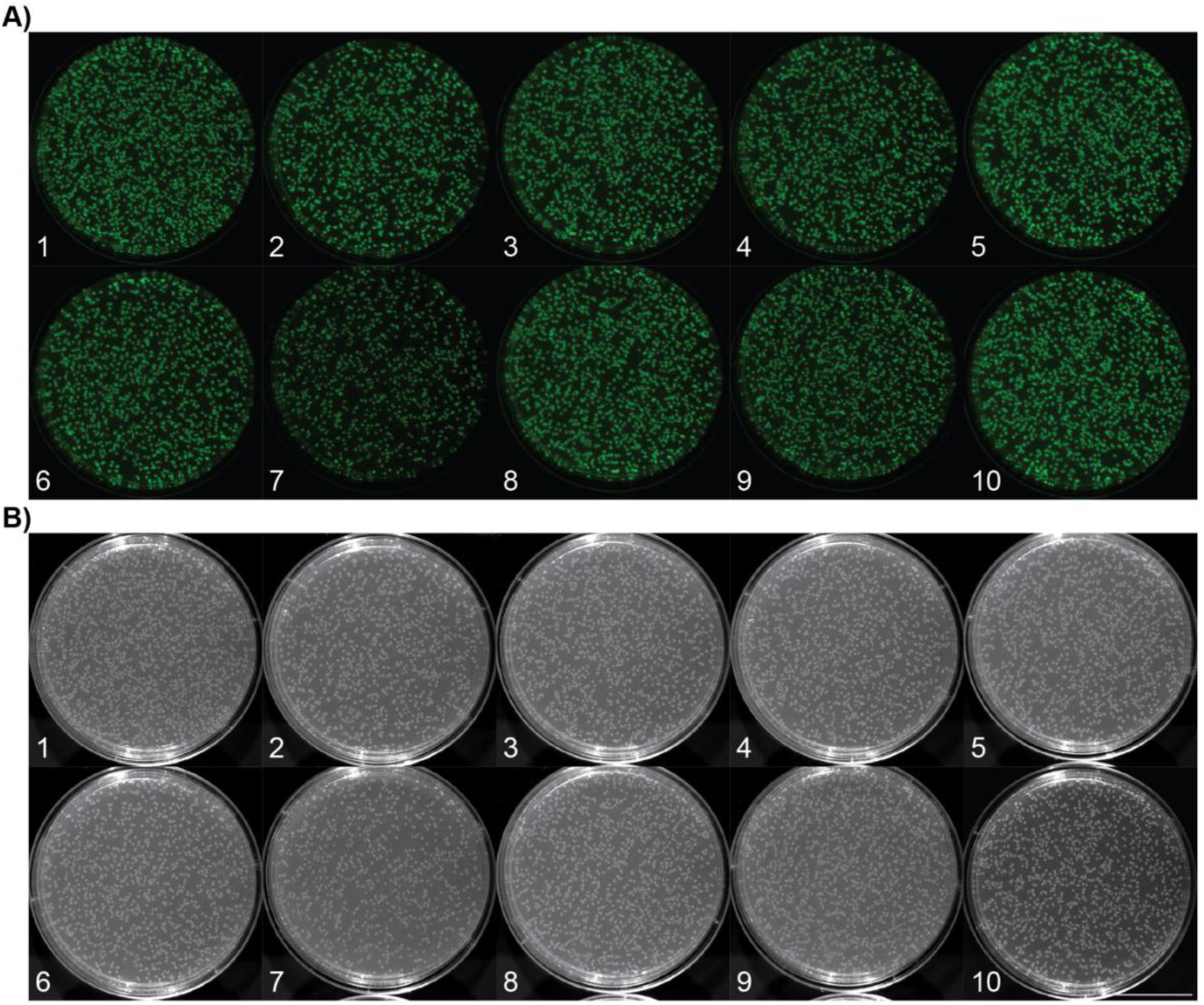
Episome retention in absence of antibiotic selection. **a)** False-colored fluorescence (GFP; excitation: 455-485 nm, emission: 508-557 nm, exposure time: 15 msec) and **b)** brightfield images of CNP^folA,TetR::gfp^ plated onto LB agar at transfers during serial culturing in LB without any antibiotic supplementation. Numerals within the panels indicate transfer numbers. Cultures reached comparable densities of OD_600_ ≍4 at each transfer. Scale bar: 30 mm.

### Unintended side effects

All relocated genomic loci were mutation-free. Yet, some of the relocation mutants acquired background mutations in their native chromosome while undergoing *in vivo* cut’n’paste (Supp. Table 7). In general, it remains unclear whether these background mutations have any functional relevance or are merely coincidental genetic drift.

## Discussion

We developed *in vivo* cut’n’paste - an easy-to-use procedure to relocate chromosomal loci onto an episome in *E. coli* that does not rely on any *in vitro* DNA manipulations. We successfully relocated genomic loci encoding metabolic and essential genes, generated a genomic alteration >100 kb, and performed up to three consecutive edits in a single strain. All relocated loci, including the 140 kb *macB*-*nadA* relocation, were mutation-free. Due to its inherent robustness, as each editing step is accompanied by positive antibiotic selection, *in vivo* cut’n’paste is well-suited for implementation in non-specialist laboratories or via unsupervised lab automation.

*In vivo* cut’n’paste can be leveraged to precisely characterise gene function. One round of *in vivo* cut’n’paste effectively creates a complementation mutant in which the gene of interest is relocated onto a single-copy extra-chromosomal replicon. Thus, *in vivo* cut’n’paste inherently overcomes the limitations of traditional sequential “delete and complement” procedures hampering the investigation of essential genes while also circumventing any convoluting copy number effects present when using traditional multi-copy plasmid backbones.

By iterating the *in vivo* cut’n’paste procedure, arbitrary modules of genes, like those obtained here (Fig. 3 & 4), can be created. For example, genes that are co-modulated or functionally connected but natively dispersed across the chromosome can easily be clustered onto a single extra-chromosomal replicon^36^. Utilising the origin of transfer (oriT) encoded on the episome, such modules can be transferred to other strains or species via conjugation without the size-limitations imposed by transformation^37^, and, hence, potential future adaptations of *in vivo* cut’n’paste to other bacterial species will only broaden its utility (see Supplementary Text).

Its ability to relocate genomic loci makes *in vivo* cut’n’paste particularly well-suited for studying how genome architecture impacts cell functioning, a topic of which we are only beginning to scratch the surface^38,39^. Ultimately, *in vivo* cut’n’paste can facilitate alternative avenues to obtain minimal genomes via bottom-up approaches to complement traditional, ineffectual top-down genome reduction attempts.

Moreover, relocations of essential genes, such as demonstrated here in CNP^folA^, are highly desirable in the context of large-scale fermentation applications in green biotechnology as they allow to stably maintain transferable extra-chromosomal replicons in production strains without incurring the metabolic costs and legal requirements associated with continual antibiotic selection^40^.

Akin to the graphical user interface in personal computing^1^, *in vivo* cut’n’paste does not enable any feature that is not already present in existing techniques. We also do not claim that *in vivo* cut’n’paste is faster or more efficient compared to other large-scale genome editing procedures. Instead, the main contribution of *in vivo* cut’n’paste lies in its ease of use, as it does not require any DNA manipulations outside of the bacterial cell, not even for the engineering of helper plasmids. While specialists in molecular biology may view multi-fragment DNA assemblies and PCRs of large, and potentially complex, DNA pieces as trivial undertakings, such tasks can appear as insurmountable hurdles to researchers utilising bacteria in fields such as biophysics, protein structure characterisation, evolutionary biology, or large-scale fermentation. *In vivo* cut’n’paste is an attempt to democratise large-scale genome editing and, thus, to open up the world of large-scale genome editing to non-specialists in molecular biology. In essence, *in vivo* cut’n’paste solely requires (1) to decide which genomic locus to relocate while identifying the homology regions HR^A^-HR^D^ (see Fig. 1B), (2) to complete the annotated pre-loading fragment genbank file templates (supplied in the Supplementary Materials), (3) to commercially obtain standardised synthetic DNA fragments, (4) to transform these synthetic DNA fragments and the easily curable helper plasmids into the strain of interest while culturing in appropriate selective conditions, (5) to cure helper plasmids, and, finally, (6) to send the mutants for whole genome re-sequencing. Thus, *in vivo* cut’n’paste empowers experimenters to refactor genomes, akin to the seven genomic edits demonstrated here, without requiring specialized knowledge in molecular biology methodologies.

## Supporting information

Episome maps

Helper plasmid maps

Synthetic DNA fragment maps

Scripts

## Materials and Methods

See “Methods” document.

## Data availability

All data is publicly available (see Supp. Table 8 for details). All custom-written scripts are supplied in the Supplementary Materials. Sequencing data was deposited at NCBI (PRJNA1271273; see Supp. Table 1, 2, and 3 for assession numbers). Sequence-verified, annotated maps of episomes and helper plasmids are supplied in the Supplementary Materials. All *in vivo* cut’n’paste homology regions are listed in Supp. Table 6. Strains are available for non-commercial use through the Belgian Coordinated Collections of Microorganisms (https://bccm.belspo.be; see Supp. Table 1 for assession numbers). Helper plasmids are available for non-commercial use through Addgene (https://addgene.org; see Supp. Table 3 for assession numbers).

## Authors’ contributions

SG, ABB, and MHHN developed the initial concept of *in vivo* cut’n’paste. SG implemented, tested, and refined *in vivo* cut’n’paste iteratively. SG performed all experiments, analysed the data, and wrote the manuscript. All authors read the final version of the manuscript.

## Acknowledgement and funding information

The authors would like to acknowledge Valeria Stavila (The Novo Nordisk Foundation Center for Biosustainability, Technical University of Denmark, Kongens Lyngby, Denmark) for technical assistance. S.G. would like to acknowledge Alex T. Nielsen (The Novo Nordisk Foundation Center for Biosustainability, Technical University of Denmark, Kongens Lyngby, Denmark) for hosting the final period of research activity in his laboratory. *E. coli* K-12 MG1655 WT was a kind gift of Tom Shimizu (AMOLF, Amsterdam, The Netherlands). The title and the abstract of this manuscript pay homage to the article “One-step inactivation of chromosomal genes in *Escherichia coli* K-12 using PCR products”. This work was supported by The Novo Nordisk Foundation.

## Supplementary Materials

**Supplementary Figure 1:**
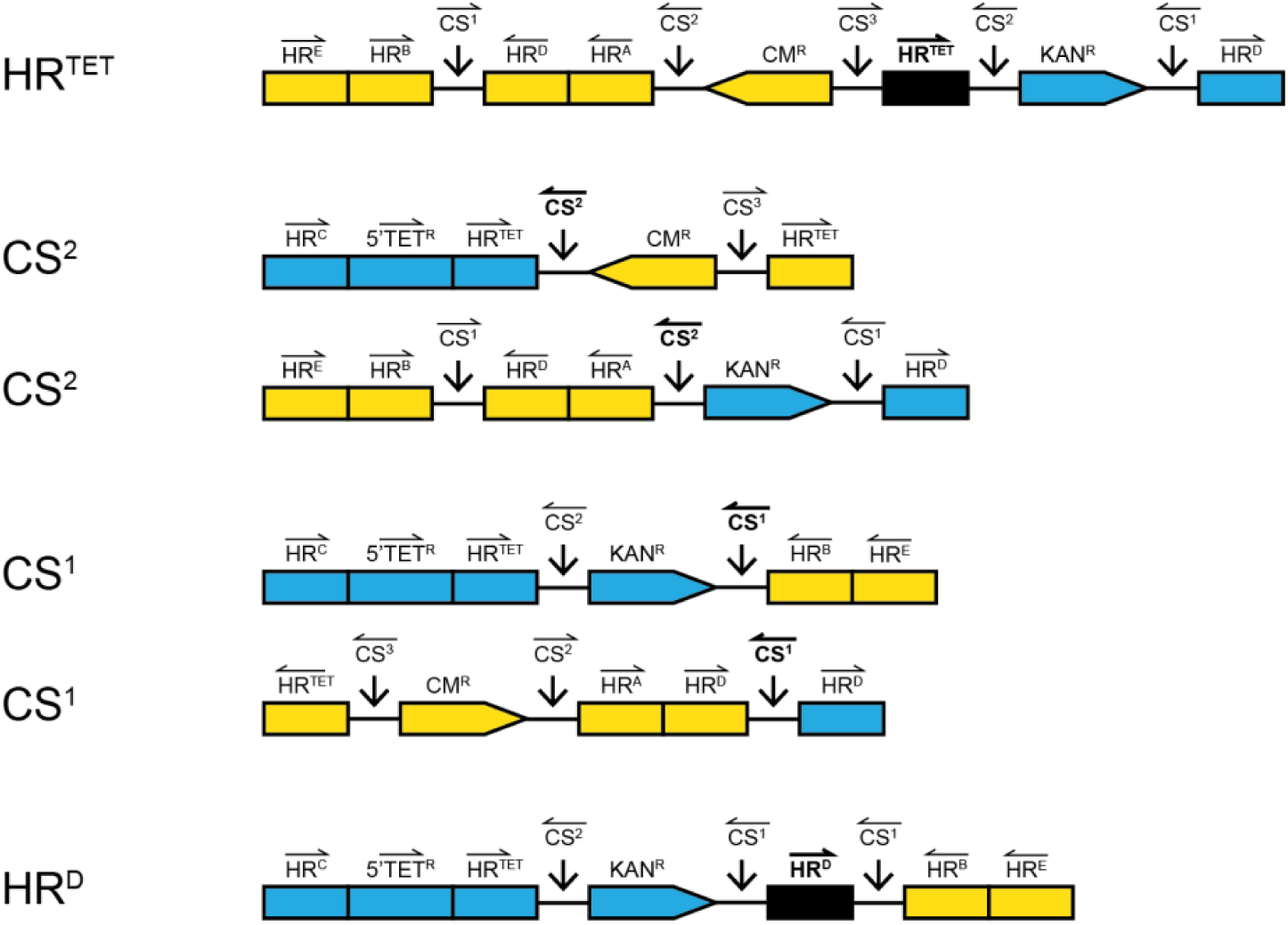
Potential for “merging” of synthetic DNA fragments during simultaneous pre-loading. Illustration of “merged” dimers of the synthetic DNA fragments PRE^epi^ (yellow) and PRE^chr^ (blue) due to sequence homology at shared features. “Merging” of PRE^epi^ and PRE^chr^ causes feature loss. Locations of “merging” are highlighted in bold and, additionally, stated on the left of the illustrations. Definitions as defined in Fig. 1. Not to scale.

**Supplementary Figure 2:**
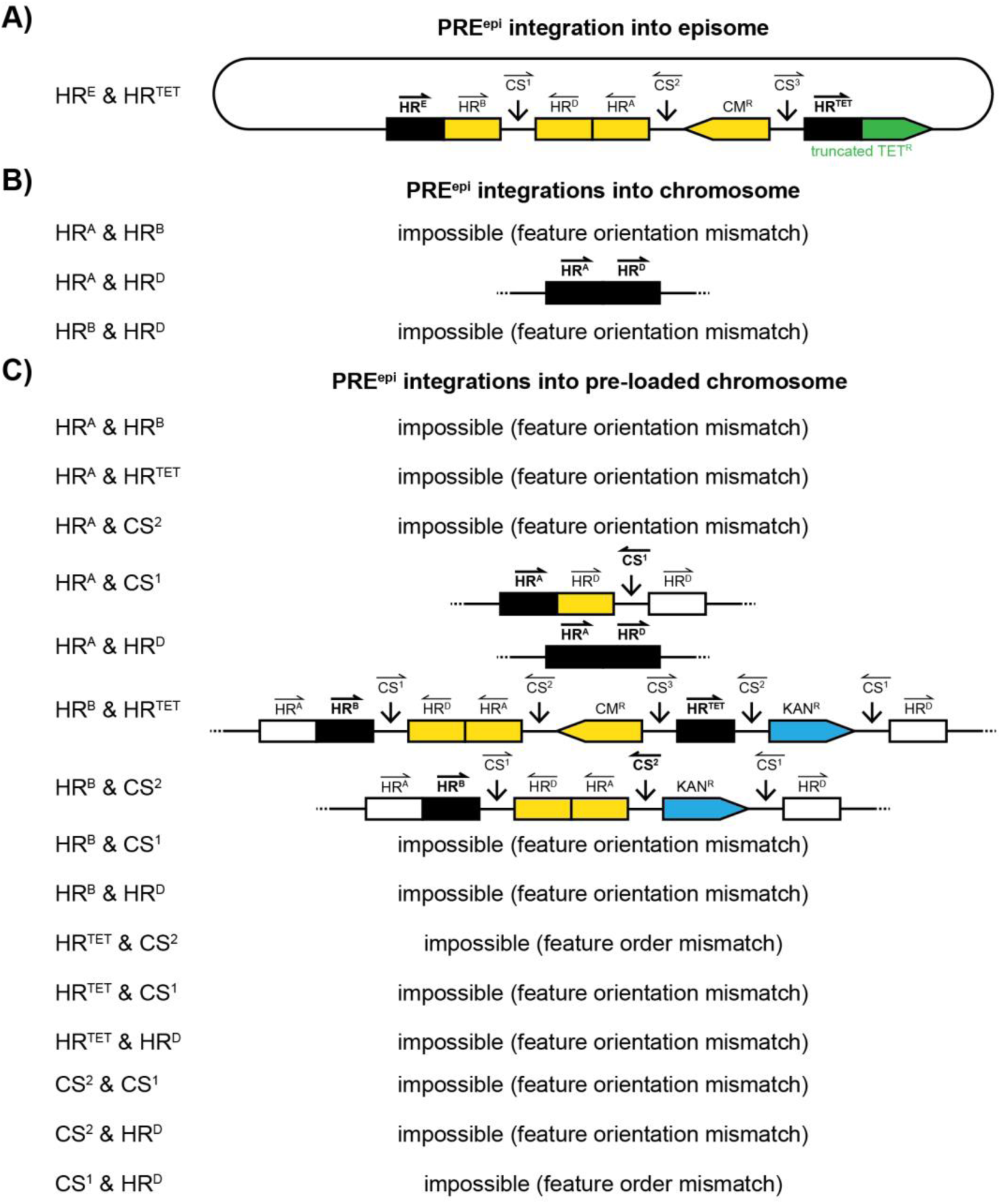
Potential integration events of synthetic DNA fragment PRE^epi^ during sequential pre-loadin. Illustration of integration events of the synthetic DNA fragment PRE^epi^ (yellow) due to homology at pairs of shared features into **a)** the episome, **b)** the native chromosome, or **c)** a native chromosome containing PRE^chr^ (blue). Integration events may cause feature loss. Many pairs of shared features cannot undergo successful integration due to their relative orientations (feature orientation mismatch) or relative locations (feature order mismatch). Integration homologies are highlighted in bold and, additionally, stated on the left side of the illustrations. Definitions as defined in Fig. 1. Not to scale.

**Supplementary Figure 3:**
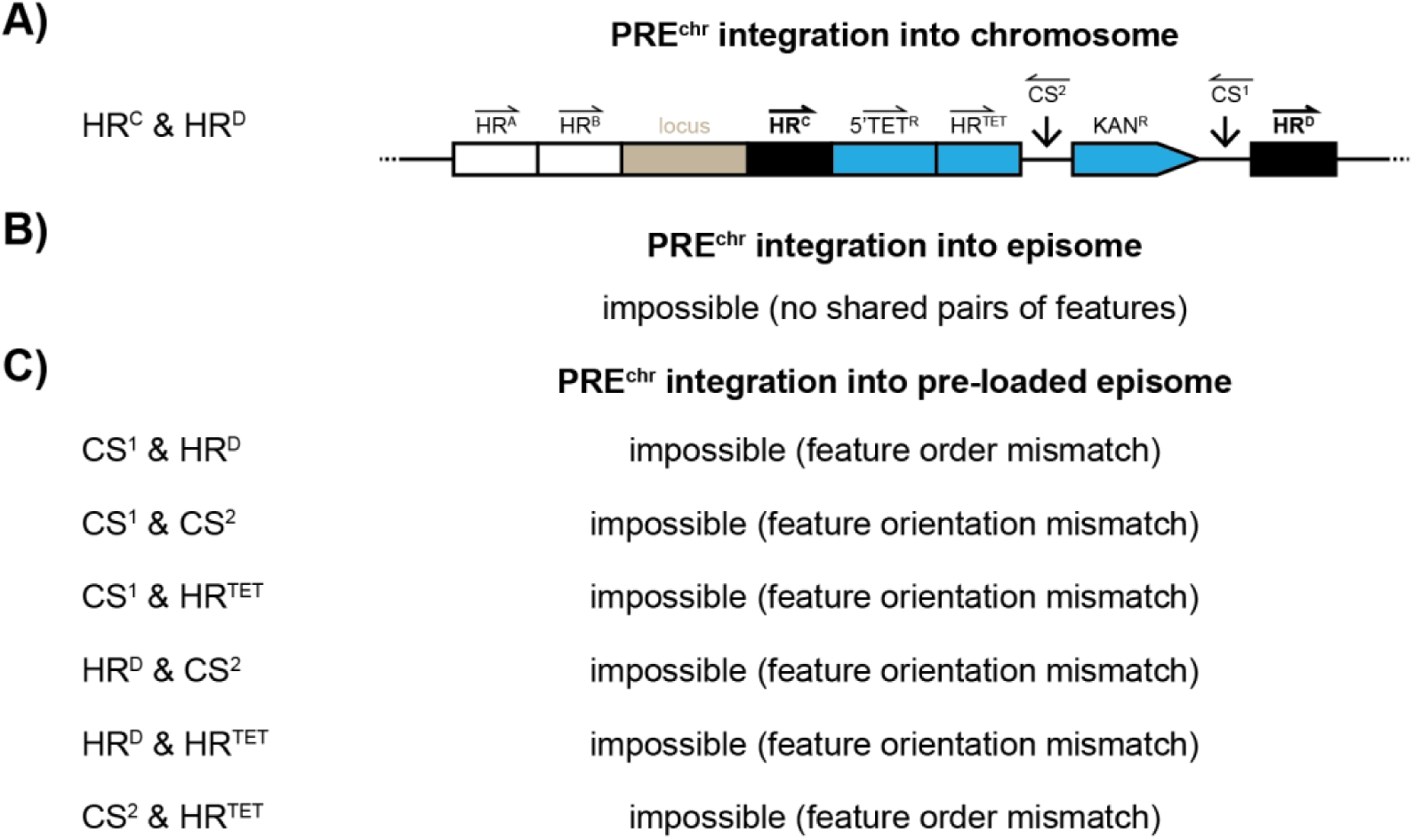
Potential integration of synthetic DNA fragment PRE^chr^ during sequential pre-loading. Illustration of integration events of the synthetic DNA fragment PRE^chr^ (blue) due to homology at pairs of shared features into **a)** the native chromosome, **b)** the episome, or **c)** an episome containing PRE^epi^. Most pairs of shared features cannot undergo successful integration due to their relative orientations (feature orientation mismatch) or relative locations (feature order mismatch). Homologies used for integration are highlighted in bold and, additionally, stated on the left side of the illustrations. Definitions as defined in Fig. 1. Not to scale.

**Supplementary Figure 4:**
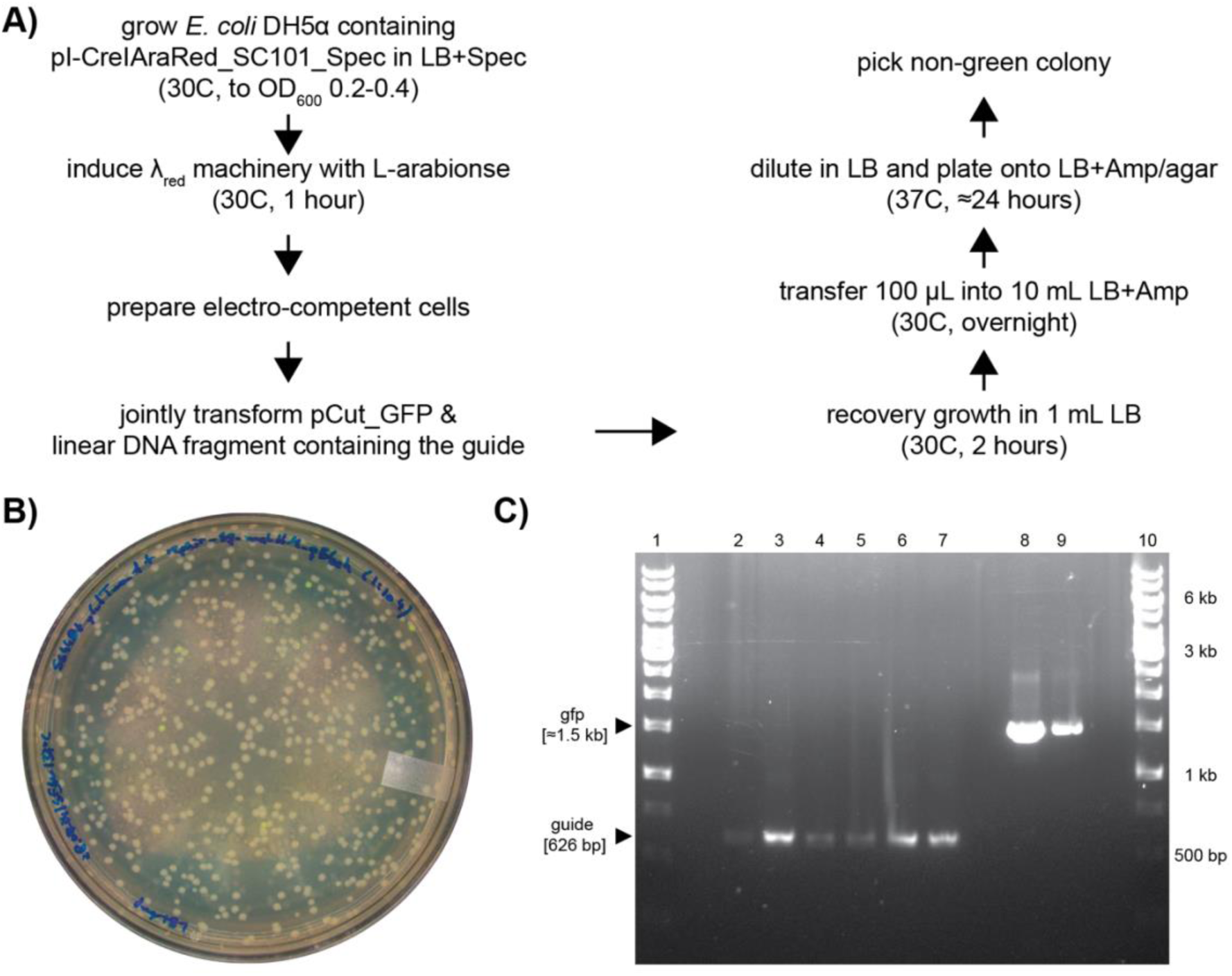
Insertion of locus-specific guide sequence into pCut_GFP via I-CreI cutting counterselection-aided plasmid recombineering. **a)** Overview of I-CreI cutting counterselection-aided plasmid recombineering workflow. **b)** Image of transformants (1:10^4^ dilution) plated onto LB-agar containing ampicillin [100 mg/L] (plate diameter: 90 mm). **c)** Agarose gel electrophoresis of diagnostic PCRs across the engineering location in pCut (primers: diag_guide_A, diag_guide_B). Non-green transformants (lanes 2-7), green transformants (lanes 8 and 9), and “gene ruler [1 kb]” DNA ladder (lanes 1 and 10). Expected band sizes are indicated by black triangles.

**Supplementary Figure 5:**
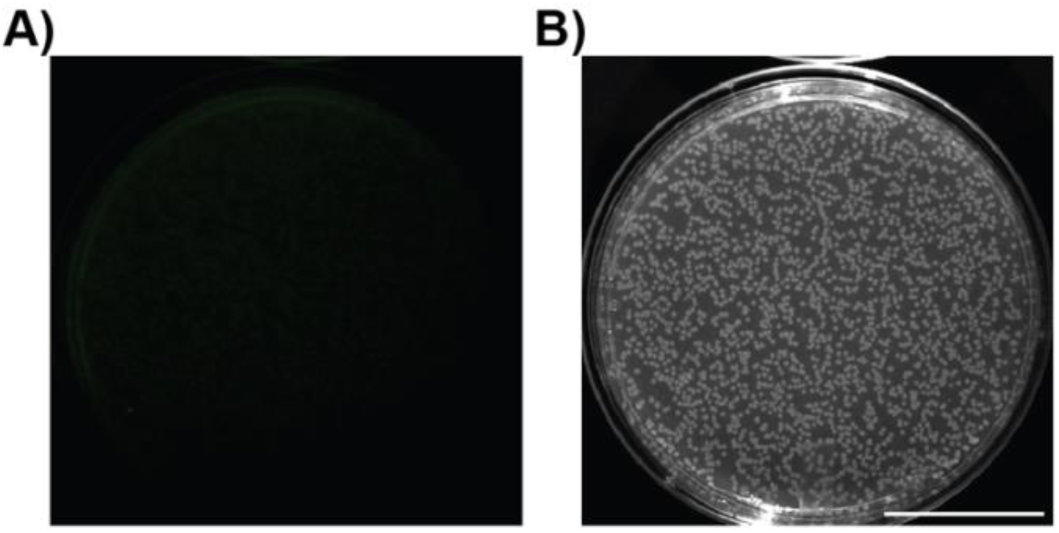
*E. coli* K-12 MG1655 WT plated on LB agar. **a)** False-colored fluorescence (GFP; excitation: 455-485 nm, emission: 508-557 nm, exposure time: 15 msec) and **b)** brightfield images of *E. coli* K-12 MG1655 WT plated onto LB agar. Identical contrast settings as in Fig. 5. Scale bar: 30 mm.

## References

1. Abbate, J. Getting small: a short history of the personal computer. Proc. IEEE 87, 1695–1698 (1999).

2. Dubois, P. F. Guest Editor’s Introduction: Python: Batteries Included. Comput. Sci. Eng. 9, 7–9 (2007).

3. Datsenko, K. A. & Wanner, B. L. One-step inactivation of chromosomal genes in *Escherichia coli* K-12 using PCR products. Proc. Natl. Acad. Sci. 97, 6640–6645 (2000).

4. Baba, T. et al. Construction of *Escherichia coli* K-12 in-frame, single-gene knockout mutants: the Keio collection. Mol. Syst. Biol. 2, 2006.0008 (2006).

5. Chapman, M. R. et al. Role of *Escherichia coli* Curli Operons in Directing Amyloid Fiber Formation. Science 295, 851–855 (2002).

6. Smith, K. D. et al. Toll-like receptor 5 recognizes a conserved site on flagellin required for protofilament formation and bacterial motility. Nat. Immunol. 4, 1247–1253 (2003).

7. Traxler, M. F. et al. The global, ppGpp-mediated stringent response to amino acid starvation in *Escherichia coli*. Mol. Microbiol. 68, 1128–1148 (2008).

8. Klumpp, S., Zhang, Z. & Hwa, T. Growth Rate-Dependent Global Effects on Gene Expression in Bacteria. Cell 139, 1366–1375 (2009).

9. Zhou, L., Lei, X.-H., Bochner, B. R. & Wanner, B. L. Phenotype MicroArray Analysis of *Escherichia coli* K-12 Mutants with Deletions of All Two-Component Systems. J. Bacteriol. 185, 4956–4972 (2003).

10. Liu, J., Gefen, O., Ronin, I., Bar-Meir, M. & Balaban, N. Q. Effect of tolerance on the evolution of antibiotic resistance under drug combinations. Science 367, 200– 204 (2020).

11. Lau, I. F. et al. Spatial and temporal organization of replicating *Escherichia coli* chromosomes. Mol. Microbiol. 49, 731–743 (2003).

12. Kolisnychenko, V. et al. Engineering a Reduced *Escherichia coli* Genome. Genome Res. 12, 640–647 (2002).

13. Meyer, A. J., Segall-Shapiro, T. H., Glassey, E., Zhang, J. & Voigt, C. A. Escherichia coli “Marionette” strains with 12 highly optimized small-molecule sensors. Nat. Chem. Biol. 15, 196–204 (2019).

14. Lee, K. H., Park, J. H., Kim, T. Y., Kim, H. U. & Lee, S. Y. Systems metabolic engineering of *Escherichia coli* for L -threonine production. Mol. Syst. Biol. 3, 149 (2007).

15. Atsumi, S. et al. Metabolic engineering of Escherichia coli for 1-butanol production. Metab. Eng. 10, 305–311 (2008).

16. Winans, S. C., Elledge, S. J., Krueger, J. H. & Walker, G. C. Site-directed insertion and deletion mutagenesis with cloned fragments in Escherichia coli. J. Bacteriol. 161, 1219–1221 (1985).

17. Hamilton, C. M., Aldea, M., Washburn, B. K., Babitzke, P. & Kushner, S. R. New method for generating deletions and gene replacements in Escherichia coli. J. Bacteriol. 171, 4617–4622 (1989).

18. Russell, C. B., Thaler, D. S. & Dahlquist, F. W. Chromosomal Transformation of Escherichia coli recD Strains with Linearized Plasmidst. J BACTERIOL 171, (1989).

19. Metcalf, W. W. et al. Conditionally Replicative and Conjugative Plasmids Carrying lacZa for Cloning, Mutagenesis, and Allele Replacement in Bacteria. Plasmid 35, 1–13 (1996).

20. Link, A. J., Phillips, D. & Church, G. M. Methods for generating precise deletions and insertions in the genome of wild-type Escherichia coli: application to open reading frame characterization. J. Bacteriol. 179, 6228–6237 (1997).

21. Kato, C., Ohmiya, R. & Mizuno, T. A Rapid Method for Disrupting Genes in the Escherichia coli Genome. Biosci. Biotechnol. Biochem. 62, 1826–1829 (1998).

22. Zhang, Y., Buchholz, F., Muyrers, J. P. P. & Stewart, A. F. A new logic for DNA engineering using recombination in Escherichia coli. Nat. Genet. 20, 123–128 (1998).

23. Posfai, G., Kolisnychenko, V., Bereczki, Z. & Blattner, F. R. Markerless gene replacement in Escherichia coli stimulated by a double-strand break in the chromosome. Nucleic Acids Res. 27, 4409–4415 (1999).

24. Jinek, M. et al. A Programmable Dual-RNA–Guided DNA Endonuclease in Adaptive Bacterial Immunity. Science 337, 816–821 (2012).

25. Jiang, W., Bikard, D., Cox, D., Zhang, F. & Marraffini, L. A. RNA-guided editing of bacterial genomes using CRISPR-Cas systems. Nat. Biotechnol. 31, 233–239 (2013).

26. Pyne, M. E., Moo-Young, M., Chung, D. A. & Chou, C. P. Coupling the CRISPR/Cas9 System with Lambda Red Recombineering Enables Simplified Chromosomal Gene Replacement in Escherichia coli. Appl. Environ. Microbiol. 81, 5103–5114 (2015).

27. Jiang, Y. et al. Multigene Editing in the Escherichia coli Genome via the CRISPR-Cas9 System. Appl. Environ. Microbiol. 81, 2506–2514 (2015).

28. Wang, K. et al. Defining synonymous codon compression schemes by genome recoding. Nature 539, 59–64 (2016).

29. Zürcher, J. F. et al. Continuous synthesis of E. coli genome sections and Mb-scale human DNA assembly. Nature 619, 555–562 (2023).

30. Wang, K., De La Torre, D., Robertson, W. E. & Chin, J. W. Programmed chromosome fission and fusion enable precise large-scale genome rearrangement and assembly. Science 365, 922–926 (2019).

31. Su, J. et al. A CRISPR-based chromosomal-separation technique for Escherichia coli. Microb. Cell Factories 21, 235 (2022).

32. Teufel, M., Klein, C. A., Mager, M. & Sobetzko, P. A multifunctional system for genome editing and large-scale interspecies gene transfer. Nat. Commun. 13, 3430 (2022).

33. Kondoh, H., Ball, C. B. & Adler, J. Identification of a methyl-accepting chemotaxis protein for the ribose and galactose chemoreceptors of Escherichia coli. Proc. Natl. Acad. Sci. 76, 260–264 (1979).

34. Adler, J. Chemotaxis in Bacteria. Science 153, 708–716 (1966).

35. Zhulin, I. B., Rowsell, E. H., Johnson, M. S. & Taylor, B. L. Glycerol elicits energy taxis of Escherichia coli and Salmonella typhimurium. J. Bacteriol. 179, 3196–3201 (1997).

36. Tan, J. et al. Independent component analysis of E. coli’s transcriptome reveals the cellular processes that respond to heterologous gene expression. Metab. Eng. 61, 360–368 (2020).

37. Hanahan, D. Studies on transformation of Escherichia coli with plasmids. J. Mol. Biol. 166, 557–580 (1983).

38. El Houdaigui, B., et al. Bacterial genome architecture shapes global transcriptional regulation by DNA supercoiling. Nucleic Acids Res. 47, 5648–5657 (2019).

39. Dame, R. T., Rashid, F.-Z. M. & Grainger, D. C. Chromosome organization in bacteria: mechanistic insights into genome structure and function. Nat. Rev. Genet. 21, 227–242 (2020).

40. Sevillano, L., Díaz, M. & Santamaría, R. I. Development of an antibiotic marker-free platform for heterologous protein production in Streptomyces. Microb. Cell Factories 16, 164 (2017).

