## Supplementary material for "Iterative *in vivo* cut’n’paste of chromosomal loci in *Escherichia coli* K-12 using synthetic DNA": Methods

#### Media and strains

Lysogeny broth (LB; tryptone [10 g/L], yeast extract [5 g/L], NaCl [10 g/L]). Tryptone both (TB; tryptone [10 g/L], NaCl [5 g/L]). Maltose-MacConkey agar (maltose [1% w/v], peptone [17 g/L], proteose peptone [3 g/L], bile salts no. 3 [1.5 g/L], NaCl [5 g/L], agar [13.5 g/L], Neutral Red [30 mg/L], Crystal Violet [1 mg/L]). M9 minimal medium (M9; $CaCl_2$ [1 mM], $MgSO_4$ [20 mM], M9 salts [1x], trace elements [1x], thiamine [5 mg/L]). M9 salts ($Na_2PO_4$ [6.8 g/L], $KH_2PO_4$ [3 g/L], $NH_4Cl$ [1 g/L], NaCl [500 mg/L]). Trace elements ($FeCl_3$ x6$H_2O$ [5 mg/L], $ZnSO_4$ x7$H_2O$ [1 mg/L], $CuCl_2$ x2$H_2O$ [200 μg/L], $MnSO_4$ x$H_2O$ [500 μg/L], $CoCl_2$ x6$H_2O$ [300 μg/L], $Na_2$EDTA [800 nM, pH 8.0]). Media were supplemented with antibiotics, effectors, and carbon sources as indicated.

Strains are listed in Supp. Table 1.

#### DNA transformation

Electro-competent bacteria were prepared as follows. Bacterial strains were cultivated in LB supplemented with appropriate antibiotics while shaking (250 rpm) at

37 °C (30 °C, if SC101 origin of replication was present) until approximately $OD_{600}$ 0.5 (until approximately $OD_{600}$ 0.3 when L-arabinose [0.5% w/v] was added and cultivation was continued while shaking (250 rpm) at 30 °C for another hour, if $\lambda_{red}$ machinery was induced). Bacteria were placed on ice for 20 minutes. Bacteria were spun down (11000 x g, 5 minutes, 4 °C), resuspended in 1 mL glycerol [15% v/v], and transferred to 2 mL Eppendorf tubes. Bacteria were spun down (11000 x g, 1 minute, 4 °C) and resuspended in 1 mL glycerol [15% v/v] three to five times. After another spin (11000 x g, 1 minute, 4 °C), bacteria were resuspended in 1/10th of their original culture volume of glycerol [15% v/v] and 50 µL aliquots were prepared. Aliquoted bacteria, stored on ice, were gently mixed with DNA, incubated on ice for approximately 5 minutes, and then transferred to chilled electroporation cuvettes (Bio-Rad Gene Pulser® Cuvette, 1 mm electrode gap). Bacteria-DNA mixtures were electroporated (Bio-Rad GenePulser Xcell™, 1.8 kV), immediately resuspended in 1 mL LB, and transferred to 1.5 mL Eppendorf tubes. Bacteria were cultivated in a thermoblock while shaking (750 rpm) at 37 °C for 1 hour (30 °C for 2 hours, if SC101 origin of replication was present). Bacteria were spun down (2400 x g, 2 minutes, room temperature), resuspended in approximately 1/10th of their own spent medium, and plated onto LB agar [1.5% w/v] plates containing appropriate antibiotics.

Chemically competent bacteria were prepared according to the TSS method[1]. Briefly, bacterial strains were cultivated in LB while shaking (250 rpm) at 37 °C until approximately $OD_{600}$ 0.5. Bacteria were placed on ice for 10 minutes. Bacteria were spun down (4000 x g, 10 minutes, 4 °C), resuspended in 1/50th of their original culture volume of TSS (PEG3305 [10% v/v], glycerol [10% v/v], $MgCl_2$ [10 mM], $MgSO_4$ x7$H_2O$ [10 mM]), left on ice for 10 minutes, and 25 µL aliquots were prepared. Aliquoted bacteria were stored at -70 °C up to several months. Aliquoted bacteria were thawed on ice and gently mixed with 25 μL DNA [≈100 – 300 ng] suspended in KCM (KCl [0.1 M], $CaCl_2$ [30 mM], $MgCl_2$ [50 mM]). Bacteria-DNA mixtures were incubated on ice for 30 minutes, resuspended in 1 mL LB, and cultivated in a thermoblock while shaking (750 rpm) at 37 °C for 1 hour (30 °C for 2 hours, if SC101 origin of replication was present). Bacteria were spun down (2400 x g, 2 minutes, room temperature), resuspended in approximately $1/10^{th}$ of their own spent medium, and plated onto LB agar [1.5% w/v] plates containing appropriate antibiotics.

#### *In vivo* cut'n'paste

*In vivo* cut'n'paste was performed in three sequential steps: (i) episome pre-loading, (ii) native chromosome pre-loading, and (iii) locus relocation. All synthetic DNA fragments were commercially obtained as gBlock HiFi Gene Fragments (Integrated DNA Technologies, Inc., Iowa, USA).

For episome pre-loading, bacteria containing an episome and the pRedCas9 helper plasmid were cultivated in LB supplemented with tetracycline [3 mg/L] and spectinomycin [50 mg/L] while shaking (250 rpm) at 30 °C. Bacteria were made electro-competent while inducing the $\lambda_{red}$ machinery and the synthetic DNA fragment $PRE^{epi}$ [≈100 ng] (see Supplementary Materials for an annotated maps) was transformed via electroporation followed by recovery growth in LB and plating on LB agar [1.5% w/v] plates containing chloramphenicol [25 mg/L] as described in section "DNA transformation". After overnight cultivation at 30 °C, which typically yielded very dense colony distributions, bacteria were restreaked onto LB agar [1.5% w/v] plates containing chloramphenicol [12.5 mg/L] and spectinomycin [50 mg/L], and cultivated at 30 °C for about 24 hours. Single colonies were transferred into 2 mL LB supplemented with chloramphenicol [12.5 mg/L] and spectinomycin [50 mg/L], cultivated while shaking (250 rpm) at 30 °C overnight, and finally glycerol [25% v/v] was added to prepare frozen stocks.

For native chromosome pre-loading, bacteria containing a pre-loaded episome and the pRedCas9 helper plasmid were cultivated in LB supplemented with chloramphenicol [12.5 mg/L] and spectinomycin [50 mg/L] while shaking (250 rpm) at 30 °C. The synthetic DNA fragment $PRE^{chr}$ (see Supplementary Materials for an annotated maps) was transformed analogously as described above for $PRE^{epi}$. Bacteria were plated onto LB agar [1.5% w/v] plates containing chloramphenicol [12.5 mg/L] and kanamycin [25 mg/L]. After overnight cultivation at 30 °C, which typically yielded very dense colony distributions, bacteria were restreaked onto LB agar [1.5% w/v] plates containing chloramphenicol [12.5 mg/L], kanamycin [25 mg/L], and spectinomycin [50 mg/L], and cultivated at 30 °C for about 24 hours. Single colonies were transferred into 2 mL LB supplemented with chloramphenicol [12.5 mg/L], kanamycin [25 mg/L], and spectinomycin [50 mg/L], cultivated while shaking (250 rpm) at 30 °C overnight, and finally glycerol [25% v/v] was added to prepare frozen stocks.

For locus relocation, bacteria with a pre-loaded native chromosome containing a pre-loaded episome and the pRedCas9 helper plasmid were cultivated in LB supplemented with chloramphenicol [12.5 mg/L], kanamycin [25 mg/L], and spectinomycin [50 mg/L] while shaking (250 rpm) at 30 °C. Bacteria were made electro-competent while inducing the $\lambda_{red}$ machinery and the helper plasmid pCut [≈250 ng] containing a locus-specific CRISPR guide was transformed via electroporation as in section “DNA transformation”. Resuspended bacteria-DNA mixtures were cultivated in a thermoblock while shaking (750 rpm) at 30 °C for 1 hour.

L-arabinose [0.5% w/v] was added and cultivation was continued while shaking (750 rpm) at 30 °C for another hour. Bacteria-DNA mixtures were transferred into a final volume of 10 mL LB supplemented with ampicillin [50 mg/L] and spectinomycin [50 mg/L] for overnight cultivation while shaking (250 rpm) at 30 °C. The following day, bacteria were spun down (11000 x g, 5 minutes, room temperature), resuspended in 1 mL LB, and 100 μL of the resuspended bacterial culture was plated onto LB agar [1.5% w/v] agar plates containing tetracycline [3 mg/L] and ampicillin [50 mg/L], and cultivated at 30 °C for about 24 hours which typically yielded very dense colony distributions. Bacteria were restreaked onto LB agar [1.5% w/v] plates containing tetracycline [3 mg/L] and spectinomycin [50 mg/L], and cultivated at 30 °C for about 24 hours. Single colonies were transferred into 2 mL LB supplemented with L-rhamnose [10 mM], tetracycline [3 mg/L], and spectinomycin [50 mg/L], and cultivated while shaking (250 rpm) at 30 °C for about 24 hours to cure the pCut helper plasmid. 100 μL of bacterial culture (1:$10^5$ diluted in LB) was plated onto LB agar [1.5% w/v] plates containing tetracycline [3 mg/L] and spectinomycin [50 mg/L], and cultivated at 30 °C for about 24 hours. Loss of pCut was confirmed by patching single colonies onto LB agar [1.5% w/v] plates containing ampicillin [100 mg/L]. Typically, all tested colonies were ampicillin-sensitive indicating loss of pCut. An ampicillin-sensitive single colony was transferred into 2 mL LB supplemented with tetracycline [3 mg/L] and spectinomycin [50 mg/L], cultivated while shaking (250 rpm) at 30 °C overnight, and finally glycerol [25% v/v] was added to prepare frozen stocks.

Optionally, if desired, the temperature-sensitive helper plasmid pRedCas9 was cured by cultivation in LB supplemented with tetracycline [3 mg/L] at 37 °C for ≈8 hours. Further processing was performed as describe above for the curing of pCut but in absence of spectinomycin, at 37 °C instead of 30 °C, and patching was conducted onto LB agar [1.5% w/v] plates containing spectinomycin [50 mg/L]. Typically, all tested colonies were spectinomycin-sensitive, indicating loss of pRedCas9. Thereafter, if desired, the episome was cured by serial cultivation while shaking (250 rpm) at 37 °C. Bacteria were cultivated in 2 mL LB for ≈8 hours, 10 μL of bacterial culture was transferred into 10 mL LB and cultivated for ≈16 hours (i.e., overnight), and finally 2 μL of bacterial culture was transferred into 2 mL LB and cultivated for ≈8 hours. Further processing was performed as described above for the curing of pRedCas9, but in the absence of antibiotic selection and patching was conducted onto LB agar [1.5% w/v] plates containing tetracycline [3 mg/L]. At least 12.5% of the tested colonies were tetracycline-sensitive, indicating loss of the episome.

Helper plasmid and episome construction

Locus-specific guides were inserted into the helper plasmid pCut via Golden Gate assembly. Briefly, pCut_GFP, containing a GFP expression cassette in the location targeted for guide insertion, was extracted using the NucleoSpin® Plasmid Purification kit (Macherey-Nagel, Germany) according to kit instructions. Pairs of oligos containing locus-specific guides flanked by Golden Gate overhangs (see Supp. Table 4) were annealed in NEBuffer 4 (New England Biolabs, MA, USA). Annealed pairs of oligos and pCut_GFP were mixed and treated with NEB Golden Gate Enzyme Mix (BsmBI-v2) (New England Biolabs, MA, USA) according to manufacturer's instructions, transformed into chemically competent *E. coli* DH5α as described in section "DNA transformation", plated onto LB agar [1.5% w/v] plates containing ampicillin [100 mg/L], and cultivated at 37 °C overnight. Non-green single colonies were restreaked onto LB agar [1.5% w/v] plates containing ampicillin [100 mg/L] and cultivated at 37 °C overnight. Single colonies were transferred into 2 mL LB supplemented with ampicillin [100 mg/L], cultivated while shaking (250 rpm) at 37 °C overnight, and finally glycerol [25% v/v] was added to prepare frozen stocks. Correct insertion of guides into pCut was verified via Sanger sequencing using primer diag_guide_B.

The guide targeting the malHM locus was inserted into the helper plasmid pCut via I-CreI cutting counterselection-aided plasmid recombineering (Supp. Fig. 4). Briefly, *E. coli* DH5α containing the helper plasmid pI-CreIAraRed_SC101_Spec was made electro-competent while inducing the $\lambda_{red}$ machinery, and the linear DNA fragment insert_guide_malHM [≈200 ng] (see Supp. Table 5) and pCut_GFP [≈200 ng] were jointly transformed via electroporation as described in section "DNA transformation". Bacteria were recovered in 1 mL LB at 30 °C for 2 hours. 100 μL of bacterial culture was transferred into 10 mL LB supplemented with ampicillin [100 mg/L] and cultivated while shaking (250 rpm) at 30 °C overnight. 100 μL of bacterial culture (1:$10^4$ diluted in LB) was plated onto LB agar [1.5% w/v] plates containing ampicillin [100 mg/L] and cultivated at 37 °C for about 24 hours. The plate was imaged with an automatic imager (Reshape Biotech, Denmark). Non-green single colonies were further processed (restreaking, preparation of frozen stocks, and sequence verification) as described for the Golden Gate assemblies.

Helper plasmids pCut_GFP, pRedCas9, and pI-CreIAraRed_SC101_Spec as well episome $EPI^{empty}$ were generated via uracil excision cloning[2]. The functional elements of the constructs are listed in Supp. Table 2 and 3, respectively. All constructs were verified via long-read sequencing as described in section "Sequencing". Sequence-verified, annotated maps are supplied in the Supplementary Materials.

Gene inactivation on native chromosome

*E. coli* K-12 MG1655 Δtrg::KanR was generated via λ Red recombineering. Briefly, flanking homologous regions were attached by Phusion U Hot Start polymerases (Thermo Fisher Scientific Inc., MA, USA) to the kanamycin marker of BW25113 Δcrp::KanR[3] according to manufacturer's instructions using primers trg_KanR_A and trg_KanR_B. The PCR fragment was purified with the NucleoSpin® Gel and PCR Clean-up kit (Macherey-Nagel, Germany) according to kit instructions.

*E. coli* K-12 MG1655 WT containing the helper plasmid pSIM19[4] was made electro-competent while inducing the $\lambda_{red}$ machinery (incubation at 42 °C in a shaking water bath for 20 minutes) and the purified PCR product [≈200 ng] was transformed via electroporation followed by recovery growth in LB and plating on LB agar [1.5% w/v] plates containing kanamycin [50 mg/L] as described in section "DNA transformation". After overnight cultivation at 37 °C, bacteria were restreaked onto LB agar [1.5% w/v] plates containing kanamycin [50 mg/L] and cultivated at 37 °C overnight. Single colonies were transferred into 2 mL LB supplemented with kanamycin [50 mg/L], cultivated while shaking (250 rpm) at 37 °C overnight, and finally glycerol [25% v/v] was added to prepare frozen stocks.

Gene replacement on episome

EPI$^{folA,TetR::gfp}$ was generated via CRISPR-Cas9 cutting counterselection-assisted λ Red recombineering. Briefly, flanking homolgous regions were attached by Phusion Hot Start polymerases (Thermo Fisher Scientific Inc., MA, USA) to the gfp cassette of SEGA007[5] according to manufacturer's instructions using primers gfp_tetA_A and gfp_tetA_B. The PCR fragment was purified with the NucleoSpin® Gel and PCR Clean-up kit (Macherey-Nagel, Germany) according to kit instructions.

CNP$^{folA}$ containing the helper plasmid pRedCas9 was made electro-competent while inducing the $\lambda_{red}$ machinery, and the purified PCR product [≈100 ng] and the helper plasmid pCut [≈250 ng] containing a CRISPR guide targeting the tetracycline cassette of EPI$^{folA}$ were jointly transformed via electroporation as described in section "DNA transformation". Further processing (outgrowth growth, restreaking, patching, curing of helper plasmids, and preparation of frozen stocks) was performed as described for locus relocation in section "*In vivo* cut'n'paste" but in the absence of tetracycline selection, and patching plates were additionally inspected under blue light for green fluorescence.

### Sequencing

Short-read whole genome re-sequencing was performed externally (Eurofins Scientific SE, Germany). Briefly, bacterial strains were streaked onto LB agar [1.5% w/v] plates containing tetracycline [3 mg/L] and cultivated overnight at 37 °C. Samples were shipped at room temperature for DNA extraction and Illumina® short-read sequencing. Sequencing results were returned in fastq format and analyzed with breseq[6] (version 0.38.1 with bowtie2[7] version 2.5.2 and R[8] version 3.6) using *E. coli*

K-12 MG1655 (NC_000913.3) and a manually generated episome map (see Supplementary Materials) as references. Predicted genomic alternation resulting from *in vivo* cut’n’paste were manually added to the references. Mutations were independently confirmed via Sanger sequencing. Mutations present in the ancestral $\text{CNP}^{\text{empty}}$ strain were ignored. Plots of mapped reads were generated with a custom Python[9] (version 3.12.7) script (MappedReadsWithCoverageOfNeighborhood.py, see Supplementary Materials). Briefly, a BAM file generated by breseq[6] was loaded using pysam[10]. For each position, read pileups were accessed via pysam[10] without filtering for read quality. Only reads that covered the position and both of its direct neighbors were counted using numpy[11] (version 2.1.3) and pandas[12] (version 2.2.3). Plots were generated with Matplotlib[13] (version 3.9.2).

Long-read whole genome re-sequencing was performed externally (Plasmidsaurus Inc, CA, USA). Briefly, bacterial strains were grown in 10 mL LB containing tetracycline [3 mg/L] at 37 °C to approx. $OD_{600}$ 1. Bacteria were spun down (11000 x g, 5 min, room temperature) and resuspended in 1 mL PBS buffer. Bacteria were spun down again (11000 x g, 1 min, room temperature) and resuspended in 500 μL DNA/RNA Shield solution (Zymo Research Corporation, CA, USA). Samples were shipped at room temperature for DNA extraction and Oxford Nanopore Technologies long-read sequencing. Sequencing results were returned in fastq format and analyzed as described for short-read whole genome re-sequencing.

Long-read plasmid and episome sequencing was performed externally (Unveil Bio ApS, Denmark). Briefly, helper plasmids and episomes were extracted using the NucleoSpin® Plasmid Purification kit (Macherey-Nagel, Germany) according to kit instructions. DNA concentrations were adjusted according to service provider instructions. Samples were shipped for external analysis. At Unveil.bio, DNA was prepared through tagmentation to generate a sequencing library. High-coverage assemblies were then generated *de novo* from Oxford Nanopore Technologies long-read DNA sequences, filtered to retain high-quality reads. Assemblies in fasta and genbank format, along with position-resolved quality and depth of coverage information, were returned and manually compared to known references with SnapGene® (version 4.2.11). Additionally, filtered raw reads in fastq format were returned. Raw reads were analyzed with a custom script (LongRead_MapAndCheck.sh, see Supplementary Materials). Briefly, raw reads were mapped to known references using minimap2[14] (version 2.26). Using samtools[15] (version 1.17), the resulting SAM files were sorted, converted to BAM format, indexed, and consensus sequences were generated. Subsequently, unmapped reads were mapped using minimap2 (version 2.26) to an *E. coli* reference (NC_000913.3). Results were manually inspected with the Integrative Genomics Viewer[16] (version 2.16.1) and SnapGene® (version 4.2.11).

Sanger sequencing was performed using the Mix2Seq Kit NightXpress kit (Eurofins Scientific SE, Germany). Briefly, PCR fragments were purified with the NucleoSpin® Gel and PCR Clean-up kit (Macherey-Nagel, Germany) according to kit instructions. Purified PCR fragments were mixed with primers according to sequencing kit instructions and shipped for external analysis. Sequencing results were returned as trace files and manually compared to known references with SnapGene® (version 4.2.11).

Phenotypic assays

Motility assay

Bacteria were cultivated in TB supplemented with tetracycline [3 mg/L] (if containing an episome) while shaking (250 rpm) at 37 °C overnight. 1 mL of overnight culture was transferred to a 2 mL Eppendorf tube. Bacteria were spun down (2400 x g, 2 minutes, room temperature) and resuspended in 1 mL M9 salts three times. 2 μL of washed bacterial culture was carefully pipetted into the center (and inside the agar) of a M9 agar [0.26% w/v] plate supplemented with glycerol [0.4% v/v] and tetracycline [3 mg/L] (if bacteria contained an episome). Additional effectors, D-ribose [100 μM] or L-Serine [100 μM], were added as stated in the individual experiments. Plates were wrapped with parafilm, placed inside an automatic time-lapse imager (Reshape Biotech, Denmark) with the agar-containing side facing away from the camera, and cultivated at 37 °C for up to five days. A water bath was placed below the imager inside the incubator. Images were automatically acquired using the setting “TOP - low” every 10 minutes.

Images were evaluated with custom-written scripts (see Supplementary Materials). Briefly, image sequences were loaded into FIJI[17] (version 2.14.0/1.54f), scaled down by a factor of 0.25, and converted to 8-bit grey-scale. Difference images were computed with a temporal shift of 18 time points using the “Stack Difference” function and then a Gaussian blur ($\sigma$ = 2.5 pixels) was applied (see “Preprocess_Loop_Stack.ijm” for details). Colony edges were detected semi-automatically. The center of the expanding colony was estimated by generating a circle from three manually selected points on the circumference of the colony at a late time point in FIJI (version 2.14.0/1.54f). Then, the radius of the circle was enlarged by 15 pixels to ensure it completely encompassed the colony. A point on the circumference of the enlarged circle was randomly picked and the intensity profile along the line connecting the point and the center of the circle was extracted. The outermost location depicting an intensity of at least 2 was used as a sampling of the colony radius. The median of 100 such samplings was used a robust estimate of the colony radius. The procedure was repeated for the preceding time point using the estimated colony radius of the previously processed time point to automatically draw an encompassing circle around the colony. This procedure was further iterated backwards through time (see "MotilityBackTracker.ijm" for details). Quality of colony radius estimation was judge by visual inspection and manually corrected if needed. Colony radii as function of time were visualized with a custom Python[9] (version 3.12.3) script using Matplotlib[13] (version 3.8.4) and pandas[12] (version 2.2.2). The initial 6 hours were excluded as the colony edges were generally not clearly visible (see "ColonyExpansionPlotting.py" for details).

<u>Maltose-utilization assay</u>

Bacteria were cultivated in LB supplemented with tetracycline [3 mg/L] while shaking (250 rpm) at 37 °C overnight. 2 μL of each overnight culture was pipetted onto a common (i.e., shared) maltose-MacConkey agar plate supplemented tetracycline [3 mg/L]. The plate was wrapped with parafilm, placed inside an automatic time-lapse imager (Reshape Biotech, Denmark) with the agar-containing side facing away from the camera, and cultivated at 37 °C for 24 hours. A water bath was placed below the imager inside the incubator. Images were automatically acquired using the settings "TOP - low", "TOP - medium", and "TOP - high" every 30 minutes.

Episome retainment assay

Bacteria were cultivated on LB agar [1.5% w/v] plates at 37 °C overnight. Single colonies were transferred into 10 mL LB and cultivated while shaking (250 rpm) at 37 °C. Every 24 hours, the bacterial cultures were transferred and sampled. For transferring, 10 μL of bacterial culture was moved into 10 mL fresh LB and cultivated while shaking (250 rpm) at 37 °C for another 24 hours. Sampling was performed by plating 100 μL of the bacterial cultures (1:$10^5$ diluted in LB) onto LB agar [1.5% w/v] plates. Plates were cultivated at 30 °C for 24 hours and subsequently green fluorescence (Ex: 455-485 nm, Em: 508-557 nm, exposure time: 15 msec) and brightfield images were acquired with an iBright™ FL1000 imaging system (Thermo Fisher Scientific Inc., MA, USA). A LB agar plate containing *E. coli* K-12 MG1655 WT was imaged as a non-fluorscent reference. Incubating, transferring, and sampling was repeated for 10 transfers in total.

After the final transfer, a single colony of each biological replicate was transferred into 10 mL LB, cultivated while shaking (250 rpm) at 37 °C overnight, and episomes were extracted and prepared for long-read sequencing as described in section “Sequencing”. The sequence of the episomes of all biological replicates was mutation-free (NCBI: SRR33800123, SRR33800122, and SRR33800121) and the episomes were additionally confirmed to constitute circular extra-chromosomal elements based the presence of overlapping individual reads along their entire known sequence map.

### References


1. Yang, Y. *et al.* Escherichia coli BW25113 Competent Cells Prepared Using a Simple Chemical Method Have Unmatched Transformation and Cloning Efficiencies. *Front. Microbiol.* **13**, 838698 (2022).
2. Cavaleiro, A. M., Kim, S. H., Seppälä, S., Nielsen, M. T. & Nørholm, M. H. H. Accurate DNA Assembly and Genome Engineering with Optimized Uracil Excision Cloning. *ACS Synth. Biol.* **4**, 1042–1046 (2015).
3. Baba, T. *et al.* Construction of *Escherichia coli* K-12 in-frame, single-gene knockout mutants: the Keio collection. *Molecular Systems Biology* **2**, 2006.0008 (2006).
4. Datta, S., Costantino, N. & Court, D. L. A set of recombineering plasmids for gram-negative bacteria. *Gene* **379**, 109–115 (2006).
5. Bayer, C. N., Rennig, M., Ehrmann, A. K. & Nørholm, M. H. H. A standardized genome architecture for bacterial synthetic biology (SEGA). *Nat Commun* **12**, 5876 (2021).
6. Deatherage, D. E. & Barrick, J. E. Identification of Mutations in Laboratory-Evolved Microbes from Next-Generation Sequencing Data Using breseq. in *Engineering and Analyzing Multicellular Systems* (eds. Sun, L. & Shou, W.) vol. 1151 165–188 (Springer New York, New York, NY, 2014).
7. Langmead, B. & Salzberg, S. L. Fast gapped-read alignment with Bowtie 2. *Nat Methods* **9**, 357–359 (2012).
8. R Foundation for Statistical Computing. R: A Language and Environment for Statistical Computing, version 3.6. Available at https://www.R-project.org.

9. Python Software Foundation. Python Language Reference, version 3.12. Available at http://www.python.org.

10. pysam - a python module for reading and manipulating files in the SAM/BAM format, version 0.22.1. Available at https://github.com/pysam-developers/pysam.

11. Harris, C. R. *et al.* Array programming with NumPy. *Nature* **585**, 357–362 (2020).

12. McKinney, W. Data Structures for Statistical Computing in Python. in 56–61 (Austin, Texas, 2010). doi:10.25080/Majora-92bf1922-00a.

13. Hunter, J. D. Matplotlib: A 2D Graphics Environment. *Comput. Sci. Eng.* **9**, 90–95 (2007).

14. Li, H. Minimap2: pairwise alignment for nucleotide sequences. *Bioinformatics* **34**, 3094–3100 (2018).

15. Li, H. *et al.* The Sequence Alignment/Map format and SAMtools. *Bioinformatics* **25**, 2078–2079 (2009).

16. Robinson, J. T. *et al.* Integrative genomics viewer. *Nature Biotechnology* **29**, (2011).

17. Schindelin, J. *et al.* Fiji: an open-source platform for biological-image analysis. *Nat Methods* **9**, 676–682 (2012).
