## Supplementary Text for "Iterative *in vivo* cut’n’paste of chromosomal loci in *Escherichia coli* K-12 using synthetic DNA"

### Supplementary Materials

Supplementary Text

Design considerations of synthetic DNA fragments for pre-loading

Basic requirements of *in vivo* cut'n'paste

Supplementary Table 1 - 8

Supplementary Table 1: Bacterial strains

Supplementary Table 2: Episomes

Supplementary Table 3: Helper plasmids

Supplementary Table 4: Oligos

Supplementary Table 5: Synthetic DNA fragments

Supplementary Table 6: *In vivo* cut'n'paste homology regions

Supplementary Table 7: Background mutations in native chromosomes of CNP mutants

Supplementary Table 8: Data deposition

Supplementary Figure 1 - 5

Supplementary Figure 1: Potential for "merging" of synthetic DNA fragments during simultaneous pre-loading

Supplementary Figure 2: Potential integration events of synthetic DNA fragment PRE$^{epi}$ during sequential pre-loading

Supplementary Figure 3: Potential integration of synthetic DNA fragment PRE$^{chr}$ during sequential pre-loading

Supplementary Figure 4: Insertion of locus-specific guide sequence into pCut_GFP via I-CreI cutting counterselection-aided plasmid recombineering

Supplementary Figure 5: *E. coli* K-12 MG1655 WT plated on LB agar

Supplementary Movies

Supplementary Movie 1: Iterative *in vivo* cut'n'paste

Supplementary Files

Synthetic_DNA_fragment_maps.zip

Episome_maps.zip

Helper_plasmid_maps.zip

Scripts.zip

**Supplementary Text**

Design considerations of synthetic DNA fragments for pre-loading

The synthetic DNA fragments $PRE^{epi}$ and $PRE^{chr}$ are flanked by homology regions targeting their integration to desired locations in the episome and in the native chromosome (Fig. 1B,C). $PRE^{epi}$ and $PRE^{chr}$ contain additional sequence homology due to the presence of several shared features. The orientations and locations of features within $PRE^{epi}$ and $PRE^{chr}$ are chosen to reduce the number of possible undesirable recombination events at these shared features. Consequently, most undesirable recombination events at shared features are hindered either because they result in the loss of at least one antibiotic marker cassette or because of the features' relative orientations or locations within the synthetic DNA fragments (Supp. Fig. 1-3). Yet, unfortunately, the design of the synthetic DNA fragments $PRE^{epi}$ and $PRE^{chr}$ does not preclude all undesired recombination events.

If present simultaneously, $PRE^{epi}$ and $PRE^{chr}$ may recombine at the shared $HR^{TET}$ feature and "merge". Such a "merged" DNA fragment, containing both $CM^{R}$ and $KAN^{R}$ marker cassettes, is able to integrate into the native chromosome via $HR^{B}$ and $HR^{D}$ causing the deletion of the genomic locus of interest while the episome would remain unaltered. "Merging" of $PRE^{epi}$ and $PRE^{chr}$ can be avoided by performing pre-loading sequentially.

$PRE^{epi}$ may integrate via $HR^{B}$ and $HR^{TET}$ into a native chromosome that already contains $PRE^{chr}$, as pre-loading of the native chromosome with $PRE^{chr}$ introduce $HR^{TET}$ next to the genomic locus of interest. As a result, both $CM^{R}$ and $KAN^{R}$ marker cassettes would be present, and the genomic locus of interest would be deleted while the episome would remain unaltered. Hence, pre-loading of the episome should be performed prior to the pre-loading of the native chromosome.

The synthetic DNA fragments used for pre-loading follow, with occasional minor variations (see Supp. Table 5 for details), the designs depicted in Fig. 1C.

Basic requirements of *in vivo* cut'n'paste

*In vivo* cut'n'paste can, in principle, be implemented in any bacterial species that fulfills the following requirements:

- a functional (heterologous) CRISPR-Cas9 system
- an efficient (heterologous) homologous recombination system
- efficient transformation of linear DNA fragments
- a low-copy episome origin of replication
- two helper plasmid origins of replication
- five antibiotic markers

**Supplementary Table 1:** Bacterial strains

| name | strain | genotype | NCBI | BCCM | source |
|---|---|---|---|---|---|
| DH5α | *E. coli* DH5α | *fhuA2Δ(argF-lacZ)U169 phoA glnV44 Φ80Δ(lacZ)M15 gyrA96 recA1 relA1 endA1 thi-1 hsdR17* | - | - | New England Biolabs |
| MG1655 Δ*trg*::KanR | *E. coli* K-12 MG1655 | Δ*trg*::KanR | - | - | This study |
| CNP$^{empty}$ | *E. coli* K-12 MG1655 | *E. coli* K-12 MG1655 WT + EPI$^{empty}$ | SAMN48903719 | LMBP 14300 | This study |
| CNP$^{trg}$ | *E. coli* K-12 MG1655 | *E. coli* K-12 MG1655 Δ*trg* + EPI$^{trg}$ | SAMN48903720 | LMBP 14301 | This study |
| CNP$^{trg,aer}$ | *E. coli* K-12 MG1655 | *E. coli* K-12 MG1655 Δ*trg* Δ*aer* + EPI$^{trg,aer}$ | SAMN48903721 | LMBP 14302 | This study |
| CNP$^{trg,aer,tsr}$ | *E. coli* K-12 MG1655 | *E. coli* K-12 MG1655 Δ*trg* Δ*aer* Δ*tsr* + EPI$^{trg,aer,tsr}$ | SAMN48903722 | LMBP 14303 | This study |
| CNP$^{malHM}$ | *E. coli* K-12 MG1655 | *E. coli* K-12 MG1655 Δ(*malH-malM*) + EPI$^{malHM}$ | SAMN48903723 | LMBP 14304 | This study |
| CNP$^{malHM,malQT}$ | *E. coli* K-12 MG1655 | *E. coli* K-12 MG1655 Δ(*malH-malM*) Δ(*malQ-malT*) + EPI$^{malHM,malQT}$ | SAMN48903724 | LMBP 14305 | This study |
| CNP$^{macB\text{-}nadA}$ | *E. coli* K-12 MG1655 | *E. coli* K-12 MG1655 Δ(*macB-nadA*) + EPI$^{macB\text{-}nadA}$ | SAMN48903725, SAMN48903728 | LMBP 14306 | This study |
| CNP$^{folA}$ | *E. coli* K-12 MG1655 | *E. coli* K-12 MG1655 Δ*folA* + EPI$^{folA}$ | SAMN48903726 | LMBP 14307 | This study |
| CNP$^{folA,gfp}$ | *E. coli* K-12 MG1655 | *E. coli* K-12 MG1655 Δ*folA* + EPI$^{folA,TetR::gfp}$ | SAMN48903727 | LMBP 14308 | This study |

**Supplementary Table 2:** Episomes

| name | description | NCBI | source |
|---|---|---|---|
| EPI$^{empty}$ | empty episome | SAMN48859298 | This study |
| EPI$^{trg}$ | episome containing the relocated *trg* locus | SAMN48859299 | This study |
| EPI$^{trg,aer}$ | episome containing the relocated *trg* and *aer* loci | SAMN48859300 | This study |
| EPI$^{trg,aer,tsr}$ | episome containing the relocated *trg*, *aer*, and *tsr* loci | SAMN48859301 | This study |
| EPI$^{malHM}$ | episome containing the relocated *malHM* locus | SAMN48859302 | This study |
| EPI$^{malHM,malQT}$ | episome containing the relocated *malHM* and *malQT* loci | SAMN48859303 | This study |
| EPI$^{macB\text{-}nadA}$ | episome containing the relocated *macB-nadA* locus | see CNP$^{macB\text{-}nadA}$ | This study |
| EPI$^{folA}$ | episome containing the relocated *folA* locus | SAMN48859304 | This study |
| EPI$^{folA,TetR::gfp}$ | episome containing the relocated *folA* locus and *gfp* (in place of the tetracycline resistance cassette) | SAMN48859305 | This study |

**Supplementary Table 3:** Helper plasmids

| name | description | NCBI | Addgene ID | source |
|---|---|---|---|---|
| pRedCas9 | arabinose-inducible $\lambda_{red}$ machinery, salicylate-inducible *cas9*, temperature-sensitive SC101 origin of replication, spectinomycin marker cassette | SAMN48859295 | 240153 | This study |
| pCut_GFP | constitutively expressed *gfp*, constitutively expressed constant guide array (CS1, CS2, CS3), rhamnose-inducible “self-curing” guide array, constitutively expressed tracrRNA, I-CreI target site, two BsmBI target sites, ColA origin of replication, ampicillin marker cassette | SAMN48859296 | 240154 | This study |
| pI-CreIAraRed_SC101_Spec | arabinose-inducible $\lambda_{red}$ machinery, constitutively expressed *i-CreI* gene, temperature-sensitive SC101 origin of replication, spectinomycin marker cassette | SAMN48859297 | 240155 | This study |
| pSIM19 | temperature-inducible $\lambda_{red}$ machinery, temperature-sensitive SC101 origin of replication, spectinomycin marker cassette | - | - | Datta, 2006 |

**Supplementary Table 4:** Oligos

| name | sequence (5’→3’) | notes |
|---|---|---|
| diag_guide_A | GTATCCTCCTGGCATCTTCTAGGAC | diagnostic, pCut_GFP, insertion of locus-specific guide sequence |
| diag_guide_B | GGCCCCAAGGGGTTATGCTAG | diagnostic, pCut_GFP, insertion of locus-specific guide sequence |
| guide_trg_A | GCACCGGTGTGAAAATGTTCAAGG | Golden Gate assembly, pCut_GFP, insertion locus-specific guide sequence, *trg* locus |
| guide_trg_B | AAACCCTTGAACATTTTCACACCG | Golden Gate assembly, pCut_GFP, insertion locus-specific guide sequence, *trg* locus |
| guide_aer_A | GCACTACTGCATTAATCGGTATGC | Golden Gate assembly, pCut_GFP, insertion locus-specific guide sequence, *aer* locus |
| guide_aer_B | AAACGCATACCGATTAATGCAGTA | Golden Gate assembly, pCut_GFP, insertion locus-specific guide sequence, *aer* locus |
| guide_tsr_A | GCACGCAAAATGTTATCGGGCATA | Golden Gate assembly, pCut_GFP, insertion locus-specific guide sequence, *tsr* locus |
| guide_tsr_B | AAACTATGCCCGATAACATTTTGC | Golden Gate assembly, pCut_GFP, insertion locus-specific guide sequence, *tsr* locus |
| guide_malHM_A | GCACTCTATGATGGAAATATTAAC | Golden Gate assembly, pCut_GFP, insertion locus-specific guide sequence, *malHM* locus |
| guide_malHM_B | AAACGTTAATATTTCCATCATAGA | Golden Gate assembly, pCut_GFP, insertion locus-specific guide sequence, *malHM* locus |
| guide_malQT_A | GCACAAGTAGAGTGCGGTTGATGC | Golden Gate assembly, pCut_GFP, insertion locus-specific guide sequence, *malQT* locus |
| guide_malQT_B | AAACGCATCAACCGCACTCTACTT | Golden Gate assembly, pCut_GFP, insertion |

| | | |
|---|---|---|
| | | locus-specific guide sequence, *malQT* locus |
| guide_macBnadA_A | GCACTCTCATTGTGTACAT CCTAA | Golden Gate assembly, pCut_GFP, insertion locus-specific guide sequence, *macB-nadA* locus |
| guide_macBnadA_B | AAACTTAGGATGTACACAA TGAGA | Golden Gate assembly, pCut_GFP, insertion locus-specific guide sequence, *macB-nadA* locus |
| guide_folA_A | GCACTTATCCGGCCTTCCT ATATC | Golden Gate assembly, pCut_GFP, insertion locus-specific guide sequence, *folA* locus |
| guide_folA_B | AAACGATATAGGAAGGCCG GATAA | Golden Gate assembly, pCut_GFP, insertion locus-specific guide sequence, *folA* locus |
| guide_tet_A | GCACGGATGCTGTAGGCA TAGGCT | Golden Gate assembly, pCut_GFP, insertion locus-specific guide sequence, Tet resistance cassette |
| guide_tet_B | AAACAGCCTATGCCTACAG CATCC | Golden Gate assembly, pCut_GFP, insertion locus-specific guide sequence, Tet resistance cassette |
| trg_KanR_A | GCCGATGACTTTCTATCAG GAGTAAACCTGGACGAGA GACAACGGTAATGATTCCG GGGATCCGTCGACC | homologous recombineering, native chromosome, replace *trg* gene with Kan resistance cassette |
| trg_KanR_B | GGGATCTGTCGATCCCTC CTTGAACATTTTCACACCG TAGCGAAACTAACTGTAGG CTGGAGCTGCTTCG | homologous recombineering, native chromosome, replace *trg* gene with Kan resistance cassette |
| gfp_tetA_A | AGACCAGAAACAAAAAAA GGCCCCCCGTTAGGGAGG CCTTCAATAATTGGTCATTT GTAGAGCTCATCCATGCCA TGTG | homologous recombineering, $EPI^{folA}$, replace Tet resistance cassette by *gfp* |
| gfp_tetA_B | AATCCTGGTGTCCCTGTTG ATACCGGGAAGCCCTGGG CCAACTTTTGGCGGCGTT GCGCTTGACGGCTAG | homologous recombineering, $EPI^{folA}$, replace Tet resistance cassette by *gfp* |

**Supplementary Table 5:** Synthetic DNA fragments

| name | sequence (5'→3') | notes |
|---|---|---|
| insert_guide_malHM | CTGACAGGGCGGGGTTTTT<br>TTTTTAATTAAAGTCTGGTCC<br>TCGAGTCTGGTTATAATCCC<br>TATCAGTGATAGAGATTGAC<br>ATCCCTATCAGTGATAGAGA<br>TACTGAGCACTCTATGATGG<br>AAATATTAACGTTTTAGAGCT<br>AGAAATAGCAAGTTAAAATAA<br>GGCTAGTCCGTTATCAACTT<br>GAAAAAGTGGCACCGAGTC<br>GGTGCTTTTTTATGCAGCCT<br>ACCGGTATCCTAGGCTGCTG<br>CCACCGCTGAGCAATAACTA<br>GCATAACCCCTTGGGGCCT<br>CTAAACGGGTCTTGAGGGG<br>TTTTTTGTTTGTATGGTTTCT<br>TAGACGTCCAAATATGTATCC<br>GCTCATGAGACAATAACCCT<br>GATAAATGCTTCAATAATATT<br>GAAAAAGGAAGAGTATGAGT<br>ATTCAACATTTCCGTGTCGC<br>CCTTATTCCCTTTTTTGCGG<br>CATTTTGCCTTCCTGTTTTT<br>GCTCACCCAGAAACGCTGG<br>TGAAAGTAAAAGATGCTGA | I-CreI counterselection-aided plasmid recombineering, pCut_GFP, insertion of locus-specific guide sequence, *malHM* locus |
| PRE_epi_trg | see annotated map in Supplementary Materials | episome pre-loading, *trg* relocation |
| PRE_chr_trg | see annotated map in Supplementary Materials | native chromosome pre-loading, *trg* relocation |
| PRE_epi_aer | see annotated map in Supplementary Materials | episome pre-loading, *aer* relocation |
| PRE_chr_aer | see annotated map in Supplementary Materials | native chromosome pre-loading, *aer* relocation |
| PRE_epi_tsr | see annotated map in Supplementary Materials | episome pre-loading, *tsr* relocation |
| PRE_chr_tsr | see annotated map in Supplementary Materials | native chromosome pre-loading, *tsr* relocation |
| PRE_epi_malHM | see annotated map in Supplementary Materials | episome pre-loading, *malHM* relocation |
| PRE_chr_malHM | see annotated map in Supplementary Materials | native chromosome pre-loading, *malHM* relocation |

| | | |
|---|---|---|
| PRE_epi_malQT | see annotated map in Supplementary Materials | episome pre-loading, *malQT* relocation |
| PRE_chr_malQT | see annotated map in Supplementary Materials | native chromosome pre-loading, *malQT* relocation |
| PRE_epi_macB-nadA | see annotated map in Supplementary Materials | episome pre-loading, *macB-nadA* relocation |
| PRE_chr_macB-nadA | see annotated map in Supplementary Materials | native chromosome pre-loading, *macB-nadA* relocation |
| PRE_epi_folA | see annotated map in Supplementary Materials | episome pre-loading, *folA* relocation |
| PRE_chr_folA | see annotated map in Supplementary Materials | native chromosome pre-loading, *folA* relocation |

**Supplementary Table 6:** *In vivo* cut'n'paste homology regions

| name | sequence (5'→3') | notes |
|---|---|---|
| HR$^{TET}$ | TACTGCCGGGCCTCTTGCGGGATATCGT<br>CCATTCCGACAGCATCGCCAGT | homology region inside tetracycline marker cassette |
| HR$^{E_trg}$ | CTGTCCCTTATTCGCACCTGGCGGTGCT<br>CAACGGGAATCCTGCTCTGCGAGGCTG<br>GCCGTAAACGAGAAAAGCCAACCTGCG<br>GGTTGGCTTTTTTATGCA | homology region on episome, *trg* relocation |
| HR$^{A_trg}$ | CGCGTAGATGCGACGTTCTCTTCTGGTT<br>GCCAGTTGATGATTAACGCTATTCGAAAA<br>TCAATGCCGTTCTGAAAGGTGAAGGGAT<br>CTGTCGATCCCTCCT | outside homology region, *trg* relocation |
| HR$^{B_trg}$ | TGAACATTTTCACACCGTAGCGAAACTAA<br>CTGGTTCACCCGCTCCGCGAGGTTCTG<br>CCGACACAGAATGTTTGTGCAGACGGAA<br>TACATCCACCGCCTCA | inside homology region, *trg* relocation |
| HR$^{C_trg}$ | ACCGTTGTCTCTCGTCCAGGTTTACTCCT<br>GATAGAAAGTCATCGGCACTTGACGGTA<br>ATTACTTATGACCAAAAACCTCTTTGCGT<br>CACATTTTTACAAC | inside homology region, *trg* relocation |
| HR$^{D_trg}$ | AATTATTGCATAGAAAATTAATGCTATTGG<br>AATGAAGAGAGTAACGAATGGCTAAACT<br>GGCTAACCCGAATGACGAAAAATTCTGG<br>AAAACAGCACGCAA | outside homology region, *trg* relocation |
| HR$^{E_aer}$ | GGTAATTACTTATGACCAAAAACCTCTTT<br>GCGTCACATTTTTACAACGGGAGACCAG<br>AAACAAAAAAAGGCCCCCCGTTAGGGAG<br>GCCTTCAATAATTGG | homology region on episome, *aer* relocation |
| HR$^{A_aer}$ | GACGCGCCTTATCCGGCCTACACCCGCT<br>ACACACCCCGCAGGCCTGATAAGATGCG<br>CCAGCATCGCATCAGGCATTGTGCTCCA<br>ACCGCCGGATCCGGCA | outside homology region, *aer* relocation |
| HR$^{B_aer}$ | TACCGATTAATGCAGTACCGTCACCGCGT<br>CTTCCAGTCGGCTGGCGCGGTGTTTCAC<br>CATCGCCGACACCTGCGCACTCTCTTCC<br>ACCAGCTCGGCATTT | inside homology region, *aer* relocation |
| HR$^{C_aer}$ | CCCCGCTGCGGTTATCTTTAACCGATTAA<br>TTTGATTTAGATCGCAATTTGCGATTTAAA<br>CACAAATCTAATTCCTTGATTTAAAATACT<br>TTCACTCTGTT | inside homology region, *aer* relocation |
| HR$^{D_aer}$ | ACTATACGAAAACGTTAATTATCTTGCCCA<br>AAAATCAGGCAATTATTGCCCTGAAAACG<br>TGCATTTGCGCAGCAATCATCAAATCCAT<br>ACCCGACAAAAA | outside homology region, *aer* relocation |
| HR$^{E_tsr}$ | ACAAATCTAATTCCTTGATTTAAAATACTT<br>TCACTCTGTTAGACAAAAAAACCCGCCG<br>CAGCGGGTCTTTGAGCC | homology region on episome, *tsr* relocation |

| | | |
|---|---|---|
| $HR^{A\_tsr}$ | ACTAATCATCTGCGGTTGCGTCACCGCA<br>GCGGGGGCACTGGCGTACATCTTCCTG<br>GTTCGTCAGCCGATCAACGACCCACGTA<br>AAGATTAATCTCCTTAT | outside homology region, *tsr* relocation |
| $HR^{B\_tsr}$ | GCCCGATAACATTTTGCTTATCGGGCATT<br>TTCATGGCGATTAAAATGTTTCCCAGTTC<br>TCCTCGCTATCTGCCACGGCCATTTTACG<br>CGGCGCAGCTGGC | inside homology region, *tsr* relocation |
| $HR^{C\_tsr}$ | GCCTGGAAAGGAAAAACTTTATGAATATC<br>CCGCGGAGATTACGCCGAGTGAATTTTAT<br>TCACACTCTGAATTTAAAAAGCCATTATTT<br>ACATAAATTATT | inside homology region, *tsr* relocation |
| $HR^{D\_tsr}$ | CACAATATAAATATGTGATATGAATCACATA<br>TTTATCGTCACTTAAACGACGCCTTTGCC<br>GCTCAACCGCAAAACTGACCGCTTACAT<br>CCCTAAAATAAC | outside homology region, *tsr* relocation |
| $HR^{E\_malHM}$ | CTGTCCCTTATTCGCACCTGGCGGTGCT<br>CAACGGGAATCCTGCTCTGCGAGGCTG<br>GCCGTAAACGAGAAAAGCCAACCTGCG<br>GGTTGGCTTTTTTATGCA | homology region on episome, *malHM* relocation |
| $HR^{A\_malHM}$ | TAAAAACAATACTATTGCCGTGACTCAGA<br>GCACGAAAGAGAATTATCGTAAGTGGGA<br>AAACAAATAACGTAAAAATAATAAGCTCTA<br>TGATGGAAATATT | outside homology region, *malHM* relocation |
| $HR^{B\_malHM}$ | AACCGGCGAACGATTCAGATTGCAGACG<br>AAAGAAAAAAAGGCGCTCCGTGGAGCG<br>CCGAATAACAGTCACAAGTTGGGATAAC<br>GTAAGTTGAGGGTGCAG | inside homology region, *malHM* relocation |
| $HR^{C\_malHM}$ | CAGTGTAAAAGGCAAGGGGTAATTACGC<br>CCCACAGTGCTGATTTTGCAACAACTGG<br>TGCGTCTCCTGGCGCACCTTTTTTTATGC<br>TTCCTTCCTGGGATA | inside homology region, *malHM* relocation |
| $HR^{D\_malHM}$ | TGAGCGATTTTTTATAGTAACTCACTTCTT<br>CTTCACTAAGAATATCCATTATCTCAATGC<br>CTTATCAGAGATTCTTTTCCTTTCGCCGG<br>TAGTGTCTGGA | outside homology region, *malHM* relocation |
| $HR^{E\_malQT}$ | TGCTGATTTTGCAACAACTGGTGCGTCT<br>CCTGGCGCACCTTTTTTTATGCTTCCTTC<br>CTGGGATAAGACAAAAAAACCCGCCGCA<br>GCGGGTCTTTGAGCC | homology region on episome, *malQT* relocation |
| $HR^{A\_malQT}$ | ATAACACTGACTGCCGGATGCGGCGTGA<br>CCGCCTTATCCGGCCTACGATTCGGGAT<br>GAATTAGTAGGCCGGATAAGACGCGTCA<br>AGCATCGCATCCGGCA | outside homology region, *malQT* relocation |
| $HR^{B\_malQT}$ | TCAACCGCACTCTACTTCTTCTTCGCTGC<br>AGCTCTGCGCCGTCTGTCCAAATCCTTC<br>AGCAACTTGTTCACGCCATCATCGGCAA<br>ACATCGACTCAAGCG | inside homology region, *malQT* relocation |

| | | |
|---|---|---|
| HR$^{C_malQT}$ | CCATCGCCAGGATGCGGTACAACACGCCCAGCAATTGCTGAAGATGATGGGGTACGGCGTGTAAGTTTAGCCGGATAACGCGCCAGATCCGGCTTACATC | inside homology region, *malQT* relocation |
| HR$^{D_malQT}$ | TCTGCATCATTCAATGCTCACCCGCGTTACGCCATCTGTTTCTATCAAACTAAACCGCACCGGCAAGAAACGCTCCACCACCGCGATATTGGTCAGCAGA | outside homology region, *malQT* relocation |
| HR$^{E_macBnadA}$ | CTGTCCCTTATTCGCACCTGGCGGTGCTCAACGGGAATCCTGCTCTGCGAGGCTGGCCGTAAACGAGAAAAGCCAACCTGCGGGTTGGCTTTTTTATGCA | homology region on episome, *macB-nadA* relocation |
| HR$^{A_macBnadA}$ | GATGTCCACCAGGGGCCAAAAGGCAATCACGCCAGTGTTATTGTGCCCGTCGAAGTAGAAGCGGCAGTCGCATAGCTCTTCTGTCTCATTGTGTACATCC | outside homology region, *macB-nadA* relocation |
| HR$^{B_macBnadA}$ | TAAAGGCAAAATGCCAGCCCGATCGGCTGGCATTTTTATCTCAAAAATTACTCTCGTGCCAGAGCATCTACTGGATCCAGTCGTGCCGCATTTCGTGCGG | inside homology region, *macB-nadA* relocation |
| HR$^{C_macBnadA}$ | CCGACATTAGCGTAATATTCGCTGTTATCGGTGTAATTTGTTTTATATGCTAAACAAGATAGTCGAATGGTGGGGAAAAGTCACTCGAATTACGCCTAAT | inside homology region, *macB-nadA* relocation |
| HR$^{D_macBnadA}$ | GTGCAGGATGCGTCATTTTACGCCTAATTCCACCGACATAGAGTTGCTTGCACAGTAAGCTCTGACGATGTCATCAGGTTTACAAAGTGAAATTGAGGAT | outside homology region, *macB-nadA* relocation |
| HR$^{E_folA}$ | CTGTCCCTTATTCGCACCTGGCGGTGCTCAACGGGAATCCTGCTCTGCGAGGCTGGCCGTAAACGAGAAAAGCCAACCTGCGGGTTGGCTTTTTTATGCA | homology region on episome, *folA* relocation |
| HR$^{A_folA}$ | ATTAACCTGCCTGCGCTGGGAAGATAAACAGTATTTTGTCCAGCCGTCGAACCGGCATAAGGATTTGGGCGAAGCGGCGGCGTCTTAAACACAGCCTGAT | outside homology region, *folA* relocation |
| HR$^{B_folA}$ | ATAGGAAGGCCGGATAAGACGCGACCGGCGTCGCATCCGGCGCTAGCCGTAAATTCTATACAAAATTACCGCCGCTCCAGAATCTCAAAGCAATAGCTGT | inside homology region, *folA* relocation |
| HR$^{C_folA}$ | ATTTTATATTCTGCTGGCGAGTCCACGCTCTCTCCCTGGACTCGCCGCATTACAATGAAACAAAAACAAACAGTTAGCTGTAAAGTGTGATTTACGTCAC | inside homology region, *folA* relocation |
| HR$^{D_folA}$ | TCTTTATTAGGATGAGGGTTTCGTTTCCGGTTCATCCGCCATGTTGCCGGTATGTTTACCTTCTTCGGTTCCCTGCCAGCCATGTCGTTGAATTAATGAC | outside homology region, *folA* relocation |

**Supplementary Table 7:** Background mutations in native chromosomes of CNP mutants

| name | gene | mutation | notes |
|---|---|---|---|
| $CNP^{trg,aer,tsr}$ | *appB* | S240S (TC**C** → TC**A**) | - |
| $CNP^{trg,aer,tsr}$ | *murP* | A344A (GC**G** → GC**A**) | - |
| $CNP^{malHM}$ | *ffh* | P168A (**C**CG → **G**CG) | essential gene[1] |
| $CNP^{malHM,malQT}$ | *ffh* | P168A (**C**CG → **G**CG) | inherited from $CNP^{malHM}$ |
| $CNP^{malHM,malQT}$ | *rpnB* | large insertion at 2,423,163 (nucleotide location in NC_000913 map) | IS1 insertion (768 bp) followed by small insertion (GGACTTCTG) |
| $CNP^{folA}$ | *uspC* | large deletion from 1,979,271 - 1,980,157 (nucleotide locations in NC_000913 map) | large deletion (887 bp), also deletes intergenic region upstream of *uspC* up to IS1H |

**Supplementary Table 8:** Data deposition

| | |
|---|---|
| Sequencing data | NCBI: PRJNA1271273 |
| Fig. 2A | see Supp. Table 1 for individual assession numbers of sequencing data |
| Fig. 2B | DOI: 10.11583/DTU.29279519<br>DOI: 10.11583/DTU.29279651 |
| Fig. 2C | DOI: 10.11583/DTU.30001390<br>DOI: 10.11583/DTU.29279651 |
| Fig. 2D | DOI: 10.11583/DTU.29279552<br>DOI: 10.11583/DTU.29279693 |
| Fig. 3B-F | DOI: 10.11583/DTU.29279558<br>DOI: 10.11583/DTU.29279723 |
| Fig. 4B | DOI: 10.11583/DTU.29279576 |
| Fig. 5 | DOI: 10.11583/DTU.29279597 |
| Supp. Fig. 4B-C | DOI: 10.11583/DTU.29279609 |
| Supp. Fig. 5 | DOI: 10.11583/DTU.29279621 |
| Supp. Table 7 | see Supp. Table 1 for individual assession numbers of sequencing data |

### References

1. Baba, T. *et al.* Construction of *Escherichia coli* K-12 in-frame, single-gene knockout mutants: the Keio collection. *Molecular Systems Biology* **2**, 2006.0008 (2006).
