## Supplementary material for "Iterative *in vivo* cut’n’paste of chromosomal loci in *Escherichia coli* K-12 using synthetic DNA": Synthetic DNA fragment maps: ReadMe.txt

Synthetic DNA fragmentes were automatically annotated with pLannotate (version 1.2.2) via http://plannotate.barricklab.org using default parameters.

Additional annotations were manually added and marked by "[MA]".
