## Supplementary material for "Iterative *in vivo* cut’n’paste of chromosomal loci in *Escherichia coli* K-12 using synthetic DNA": Scripts: ColonyExpansion_Plotting.py

```
##
# ColonyExpansion_Plotting.py
##


# libraries
import pandas as pd
import matplotlib.pyplot as plt


# functions


# parameters
data_file_path = '_FILE_PATH_'

offset_slices = 18
slices_per_hour = 6

pixel_per_mm = 690 / 90


# script

# load data into DataFrame
data_df = pd.read_csv(data_file_path, sep='\t', header=5)

# convert slice number to hours
data_df['time [days]'] = (
    data_df['slice'] + offset_slices - 1) / (slices_per_hour * 24)
# delete 'slice' column
data_df.drop(labels=['slice'], axis='columns', inplace=True)
# set 'time [hours]' column as index
data_df.set_index('time [days]', drop=True, inplace=True)
# sort index
data_df.sort_index(ascending=True, inplace=True)

# convert pixel to mm
data_df = data_df / pixel_per_mm

# filter out first 6 hours
data_df = data_df[data_df.index>0.25]

# visuailze colony radius as function of time
fig, ax = plt.subplots(figsize=(2.625,1.5), dpi=300)

ax.plot(data_df.index, data_df['_STRAIN_MEDIUM_'], '-',
        c=(255/255, 36/255, 0/255), lw=1)
ax.plot(data_df.index, data_df['_STRAIN_MEDIUM_'], '-',
        c=(100/255,149/255,237/255), lw=1)
ax.plot(data_df.index, data_df['_STRAIN_MEDIUM_'], '-',
        c='k', lw=1)

# colors
# (255/255, 36/255, 0/255) = dark red
# (250/255, 128/255, 114/255) = light red
# (182/255,208/255,226/255) = light blue
# (100/255,149/255,237/255) = blue

# axes, labels, legend, ...
ax.set_xlim(-0.05,3.05)
ax.set_ylim(-0.5,29)
ax.set_xticks([0,0.5,1,1.5,2,2.5,3])
ax.set_yticks([0,10,20])
ax.tick_params(labelbottom=True, labeltop=False, labelleft=True,
               labelright=False, bottom=True, top=True, left=True, right=True)
ax.tick_params(axis='x', labelsize=9)
ax.tick_params(axis='y', labelsize=9)
ax.set_xlabel('time [d]', fontsize=9)
ax.set_ylabel('colony radius [mm]', fontsize=9)

# save figure
plt.savefig('ColonyRadiusAsFunctionOfTime.pdf')

# clean up
del fig, ax
```
