## Supplementary material for "Iterative *in vivo* cut’n’paste of chromosomal loci in *Escherichia coli* K-12 using synthetic DNA": Scripts: LongRead_MapAndCheck.sh

```
#! /bin/bash

# required arguments
# filename/path of sequencing data fastq file
# filename/path of sequencing reference (i.e., plasmid) fasta file
# filename/path of genome reference (e.g., E. coli MG1655 if plasmid was miniprepped from an E. coli strain) fasta file


# retrieve passed arguments
fastq_f=$1
ref_f=$2
ref_genome_f=$3


# get fastq filename without extension
fastq_n=${fastq_f%.*}


# map all reads to sequencing reference

# align with minimap2 using Oxford nanopore settings to sequencing reference
minimap2 -ax map-ont $ref_f $fastq_f > ${fastq_n}_aln.sam

# mapping report
echo ""
echo "Mapping report stats:"
samtools flagstat ${fastq_n}_aln.sam
echo -n "Unmapped reads: "
samtools view -c -f 4 ${fastq_n}_aln.sam
echo ""

# sort alignment with samtools
samtools sort -o ${fastq_n}_aln_sorted.sam ${fastq_n}_aln.sam

# convert sam to bam with samtools
samtools view -S -b ${fastq_n}_aln_sorted.sam > ${fastq_n}_aln_sorted.bam

# index bam with samtools
samtools index -M -b ${fastq_n}_aln_sorted.bam

# create consensus with samtools
samtools consensus -a -m simple -c 0.51 -d 3 --min-BQ 20 --min-MQ 20 --show-ins yes --show-del no ${fastq_n}_aln_sorted.bam -f fasta -o ${fastq_n}_consensus.fa

# extract unmapped reads (i.e., reads that did not map to filtering reference)
samtools fasta -f 4 ${fastq_n}_aln.sam > ${fastq_n}_unmapped.fa


# check for genomic reads (try to map previously unmapped reads to the genome reference)

# align reads with minimap2 using Oxford nanopore settings to filtering reference
minimap2 -ax map-ont $ref_genome_f ${fastq_n}_unmapped.fa > ${fastq_n}_genomic_aln.sam

# genomic reads report
echo ""
echo "Genomic reads report stats:"
samtools flagstat ${fastq_n}_genomic_aln.sam
echo -n "Unmapped reads: "
samtools view -c -f 4 ${fastq_n}_genomic_aln.sam
echo ""
```
