## Supplementary material for "Iterative *in vivo* cut’n’paste of chromosomal loci in *Escherichia coli* K-12 using synthetic DNA": Scripts: MappedReadsWithCoverageOfNeighborhood.py

```
##
#
# MappedReadsWithCoverageOfNeighborhood.py
#
##


## libraries
import pysam
import numpy as np
import pandas as pd
import matplotlib.pyplot as plt


## parameters

# import
bam_filename = '_FILE_PATH_' # indexed BAM file

# region
region_contig = '_NAME_' # name of contig
region_start = _START_ # left end of the genomic position of region; inclusive (0-indexing)
region_stop = _STOP_ # right end of the genomic position of region; exclusive (0-indexing)

# read filtering
filter_halfwidth = 1 # half width of interval around position
filter_min_covered = 1 + 2*filter_halfwidth # minimum number of positions in interval that must be covered by read
filter_min_base_quality = 0 # minimum base quality for reads to be considered

# plotting
marker_pos_01 = _POS1_ # genomic position of marker of HRD-HRC boundary; (1-indexing)
marker_pos_02 = _POS2_ # genomic position of marker of HRB-HRA boundary; (1-indexing)
marker_pos_03 = np.nan # unused; (1-indexing)

locus_start = _POS_START_ # left end of the genomic position of the locus; inclusive (1-indexing)
locus_end = _POS_END_ # right end of the genomic position of the locus; inclusive (1-indexing)

flip_x_axis = True

# export
export_prefix = '_PREFIX_'


## script

# process BAM file with pysam to compute "filtered" coverage

# read BAM file
bamfile = pysam.AlignmentFile(bam_filename, 'rb')

# initialise counter
filtered_read_counts = {}

# perform pileup in region and loop over positions
for pileupcolumn in bamfile.pileup(contig=region_contig, start=region_start, stop=region_stop, max_depth=1000000, truncate=True):
        
    # set min base quality
    pileupcolumn.set_min_base_quality(filter_min_base_quality)
    
    # initialise counter
    read_counts = 0
    
    # loop over all pileupreads at position
    for pileupread in pileupcolumn.pileups:
        
        # get positions mapped by read as numpy array
        ref_positions_np = np.array(pileupread.alignment.get_reference_positions())
        
        # count read only if it covers at least min_covered positions in interval [position-halfwidth,position+halfwidth]
        if len(ref_positions_np[(ref_positions_np>=pileupcolumn.pos-filter_halfwidth) & (ref_positions_np<=pileupcolumn.pos+filter_halfwidth)]) >= filter_min_covered:
            read_counts += 1
    
    # store read counts in dict
    filtered_read_counts[pileupcolumn.pos] = read_counts     
# clean up
del pileupcolumn, pileupread, read_counts, ref_positions_np

# close BAM file
bamfile.close()

# convert to DataFrame
filtered_read_counts_df = pd.DataFrame.from_dict(filtered_read_counts, orient='index', columns=['cov'])
filtered_read_counts_df.index.names = ['position']
# clean up
del filtered_read_counts

# convert index to 1-indexing (as used typically, e.g., by samtools)
filtered_read_counts_df.index = filtered_read_counts_df.index + 1


# visualise coverage as function of position
fig, ax = plt.subplots(figsize=(3.0,0.8), dpi=300)

# plot total coverage
if locus_start >= 1 and locus_end >= 1:
    # plot coverage outside of locus in black
    ax.plot(filtered_read_counts_df[filtered_read_counts_df.index < locus_start].index, filtered_read_counts_df[filtered_read_counts_df.index < locus_start]['cov'], '-', c='k', lw=1)
    ax.plot(filtered_read_counts_df[filtered_read_counts_df.index > locus_end].index, filtered_read_counts_df[filtered_read_counts_df.index > locus_end]['cov'], '-', c='k', lw=1)
    # plot coverage inside locus in yellow
    ax.plot(filtered_read_counts_df[(filtered_read_counts_df.index >= locus_start) & (filtered_read_counts_df.index <= locus_end)].index, filtered_read_counts_df[(filtered_read_counts_df.index >= locus_start) & (filtered_read_counts_df.index <= locus_end)]['cov'], '-', c='y', lw=1)
else:
    # plot coverage in black
    ax.plot(filtered_read_counts_df.index, filtered_read_counts_df['cov'], '-',  c='k', lw=1)

# marker_01
ax.plot(marker_pos_01,750, 'kv', ms=5)
# marker_02
ax.plot(marker_pos_02,750, 'kv', ms=5)
# marker_03
ax.plot(marker_pos_03,750, 'kv', ms=5)

# axes, labels, legend, ...
ax.set_xlim(min(filtered_read_counts_df.index),max(filtered_read_counts_df.index))
ax.set_ylim(-50,800)
ax.set_yticks([0,300,600])
ax.tick_params(labelbottom=False, labeltop=False, labelleft=True, labelright=False, bottom=False, top=False, left=True, right=True)
ax.tick_params(axis='x', labelsize=9)
ax.tick_params(axis='y', labelsize=9)
ax.set_xlabel('position', fontsize=9)
ax.set_ylabel('cov', fontsize=9)
if flip_x_axis:
    ax.invert_xaxis()
fig.tight_layout()

# save figure
plt.savefig(export_prefix + '_Filtered_Coverage.pdf')

# clean up
del fig, ax
```
