## Supplementary material for "Iterative *in vivo* cut’n’paste of chromosomal loci in *Escherichia coli* K-12 using synthetic DNA": Scripts: MotilityBackTracker.ijm

```
// MotilityBackTracker.ijm


// functions

// compute circle (center and radius) based on 3 pairs of x-y coordinates
function computeCircle(x_0, y_0, x_1, y_1, x_2, y_2) {
	
	// compute location of mid-point of p_0 and p_1
	x_mid_0_1 = (x_0 + x_1)/2.0;
	y_mid_0_1 = (y_0 + y_1)/2.0;
	// compute location of mid-point of p_1 and p_2
	x_mid_1_2 = (x_1 + x_2)/2.0;
	y_mid_1_2 = (y_1 + y_2)/2.0;
	
	// compute the orthogonal line of mid_0_1 to line defined by p_0 and p_1
	x_orth_0_1 = x_mid_0_1 + (y_1 - y_0);
	y_orth_0_1 = y_mid_0_1 - (x_1 - x_0);
	// compute the orthogonal line of mid_1_2 to line defined by p_1 and p_1
	x_orth_1_2 = x_mid_1_2 + (y_2 - y_1);
	y_orth_1_2 = y_mid_1_2 - (x_2 - x_1);
	
	// compute the intersection point of orth_0_1 and orth_1_2 (i.e., the center of the circles)
	denominator = (x_mid_0_1 - x_orth_0_1) * (y_mid_1_2 - y_orth_1_2) - (y_mid_0_1 - y_orth_0_1) * (x_mid_1_2 - x_orth_1_2);
	if(denominator!=0){
		// compute center of the circle
		x_circle = ( (x_mid_1_2 - x_orth_1_2) * (x_mid_0_1*y_orth_0_1 - y_mid_0_1*x_orth_0_1) - (x_mid_0_1 - x_orth_0_1) * (x_mid_1_2*y_orth_1_2 - y_mid_1_2*x_orth_1_2) ) / denominator;
		y_circle = ( (y_mid_1_2 - y_orth_1_2) * (x_mid_0_1*y_orth_0_1 - y_mid_0_1*x_orth_0_1) - (y_mid_0_1 - y_orth_0_1) * (x_mid_1_2*y_orth_1_2 - y_mid_1_2*x_orth_1_2) ) / denominator;
	
		// compute radius of the circle
		r_circle = Math.sqrt(Math.sqr(x_circle - x_0) + Math.sqr(y_circle - y_0));
		
		return newArray(x_circle, y_circle, r_circle);
	}
	else {
		return newArray(0,0,-1);
	}
}

// get random point on circumference of circle
function getRandomPointOnCircumference(x_center, y_center, radius) {
	// get allowed x-range inside the radius of the current circle
	x_min = Math.ceil(x_center - radius);
	x_max = Math.floor(x_center + radius);
	
	// randomly select an x-value within the circle area
	x_random = random() * (x_max - x_min) + x_min;

	// compute y-value (randomly pick sign in front of square root)
	if(random()<0.5) {
		y_random = y_center + Math.sqrt(Math.sqr(radius) - Math.sqr(x_random - x_center));
	}
	else{
		y_random = y_center - Math.sqrt(Math.sqr(radius) - Math.sqr(x_random - x_center));
	}
	// return random point on circumference of circle
	return newArray(x_random, y_random);
}

// find outermost threshold crossing along line
function findOuterThresholdCrossing(x_cen, y_cen, x_cir, y_cir, threshold, delay) {
	// create line from center to circumference
	makeLine(x_cen, y_cen, x_cir, y_cir);
	// get profile along line
	profile = getProfile();
	// "flash" lines for visual inspection
	wait(delay);
	// remove line from image again
	run('Select None');
	// locate outermost threshold crossing
	for (i=profile.length-1; i>=0; i--) {
		if(profile[i]>threshold) {
		  return i+1;
		 }
	}
	return NaN;
}

// compute median across array of numbers
function median(array){
	// sort array while dropping NaN values
	array=Array.sort(Array.deleteValue(array,NaN));
	if(array.length==0) {
		return NaN;
	}
	if (array.length%2>0.5) {
		return array[floor(array.length/2)];
	}
	else{
		return (array[array.length/2]+array[array.length/2-1])/2;
	}
}


// parameters

delta_radius = 15;
threshold = 1;
num_sampling = 100;
vis_delay = 0;


// script

run('Clear Results');

// manually create initial guess of the circle location (by selecting 3 point in the image)

// initialize location array and counter
loc = newArray();
counter = 0;
// loop until 3 points that do not fall onto one straight line are selected or 'esc' is pressed
while(true) {
	// get current cursor location
	getCursorLoc(x, y, z, flags);
	// listen for right mouse clicks
	if(flags==16) {
		// store location of right mouse click
		loc[counter] = x;
		loc[counter+1] = y;
		loc[counter+2] = z;
		counter += 3;
		wait(500); // avoid collecting the same mouse click multiple times
	}
	// check whether 3 locations (i.e., 3 x 3 = 9 coordinates) have been collected
	if(counter>=9) {
		// check whether the 3 locations are in the same slice
		if(loc[2]==loc[5] && loc[5]==loc[8]) {
			// try to compute circle
			circle_array = computeCircle(loc[0],loc[1],loc[3],loc[4],loc[6],loc[7]);
			if(circle_array[2]!=-1) {
				break;
			}
		}
		// reset location array and counter
		loc = newArray();
		counter = 0;
	}
}

// search for circles in all slides up to the current one (going backwards)
current_slice = getSliceNumber();
for (slice=current_slice;slice>=1;slice--) {
		
	// slightly enlarge initial guessed circle
	circle_array[2] += delta_radius;
		
	radii_array = newArray();
	for (i=0;i<num_sampling;i++) {
		xy_cir = getRandomPointOnCircumference(circle_array[0],circle_array[1],circle_array[2]);
		radii_array[i] = findOuterThresholdCrossing(circle_array[0], circle_array[1], xy_cir[0], xy_cir[1], threshold, vis_delay);
	}
	// compute circle radius
	radius_new = median(radii_array);
	
	// only update and show circle if it has a valid radius
	if(!isNaN(radius_new)){
		// update circle radius
		circle_array[2] = radius_new;
		
		// show updated circle
		makeOval(circle_array[0]-circle_array[2], circle_array[1]-circle_array[2], circle_array[2]*2, circle_array[2]*2);
		run('Add Selection...');
		run('Select None');
	}
	else{
		circle_array[2] -= delta_radius;	
	}
	
	// append to results table
	append_idx = nResults;
	setResult('slice', append_idx, slice);
	setResult('radius', append_idx, radius_new);
	updateResults();
			
	// stop going backwards if 1st slice is reached
	if(slice>1) {
		setSlice(slice-1);
	}
}
```
