## Supplementary material for "Iterative *in vivo* cut’n’paste of chromosomal loci in *Escherichia coli* K-12 using synthetic DNA": Scripts: Preprocess_Stack_Loop.ijm

```
// Preprocess_Stack_Loop.ijm


// parameters
para_step = 1;
para_gap = 18;
para_scale = 25.0;
para_gb_sigma = 2.5;

im_directory_prefix = '_DIR_PATH_';
im_directory_postfix = '/A1_well/RIS1_0_TL_5_preset';

im_directories = newArray(15);
im_directories[0] = '_FILE_NAME_';
im_directories[1] = '_FILE_NAME_';
im_directories[2] = '_FILE_NAME_';
im_directories[3] = '_FILE_NAME_';
im_directories[4] = '_FILE_NAME_';
im_directories[5] = '_FILE_NAME_';
im_directories[6] = '_FILE_NAME_';
im_directories[7] = '_FILE_NAME_';
im_directories[8] = '_FILE_NAME_';
im_directories[9] = '_FILE_NAME_';
im_directories[10] = '_FILE_NAME_';
im_directories[11] = '_FILE_NAME_';
im_directories[12] = '_FILE_NAME_';
im_directories[13] = '_FILE_NAME_';
im_directories[14] = '_FILE_NAME_';


// script
setBatchMode(true);
for(i=0;i<im_directories.length;i++) {
	showStatus(i+1 + '/' + im_directories.length + ' image directories');
	run('Collect Garbage');
	File.openSequence(im_directory_prefix + im_directories[i] + im_directory_postfix, 'step=' + para_step + ' scale=' + para_scale);
	original_im_id = getImageID();
	run('8-bit');
	run('Stack Difference', 'gap=' + para_gap);
	run('Gaussian Blur...', 'sigma=' + para_gb_sigma + ' stack');
	rename(im_directories[i] + '_pre-processed');
	saveAs('tiff', '/Users/segu/Desktop/' + getTitle());
	close();
	selectImage(original_im_id);
	close();
}
setBatchMode(false);
```
