## Supplementary figures and images for "Iterative *in vivo* cut’n’paste of chromosomal loci in *Escherichia coli* K-12 using synthetic DNA"

### Supplementary Movie 1

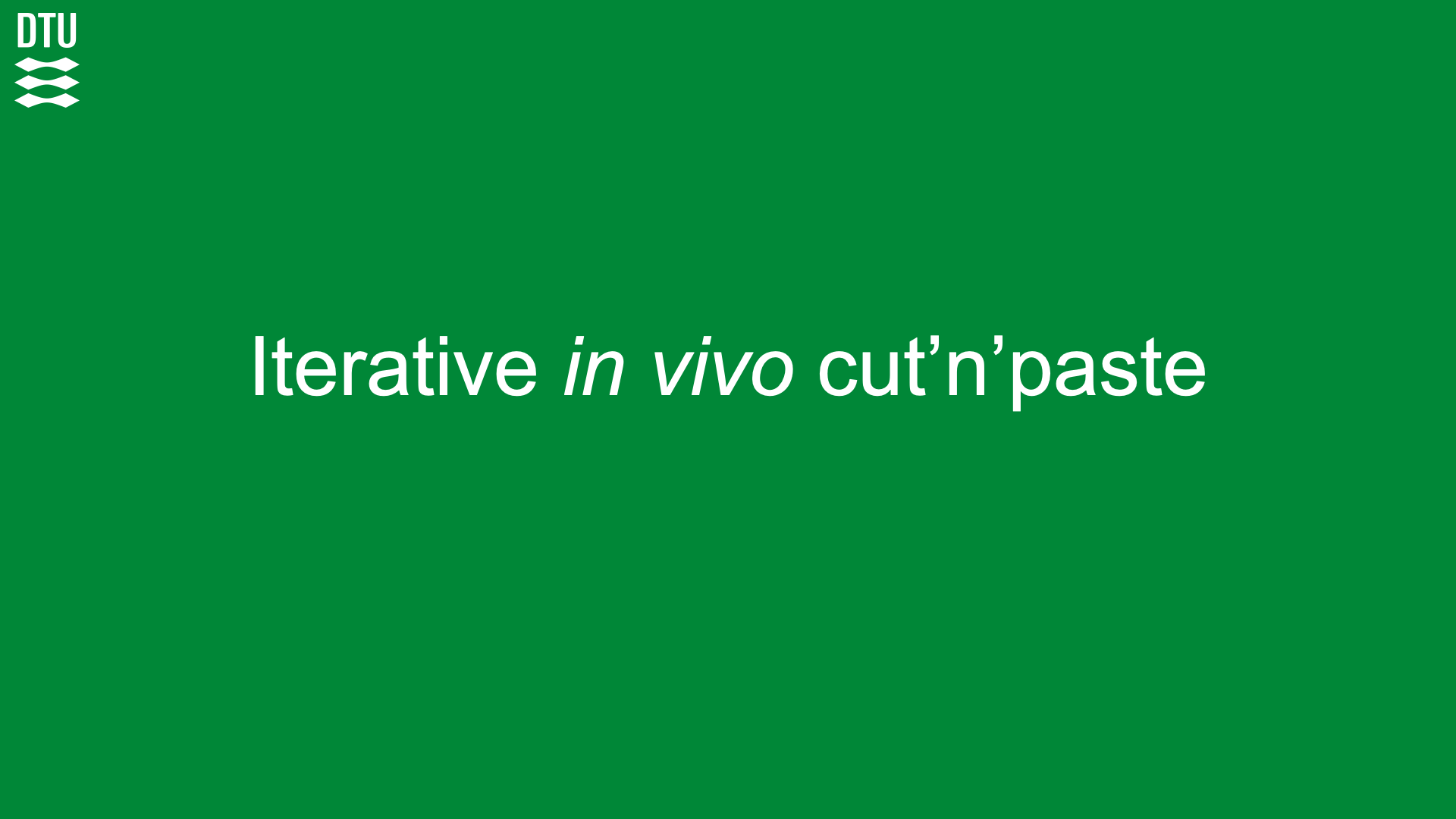
